# Landscape suitability and range expansion estimates for the North American Interior Population of trumpeter swans

**DOI:** 10.1101/2025.04.03.647047

**Authors:** Kevin W. Barnes, Thomas R. Cooper, David E. Andersen, Mike E. Estey, David W. Wolfson

## Abstract

The Interior Population of trumpeter swans (*Cygnus buccinator*) has grown and expanded substantially since reintroduction efforts began in the 1950s. To support management of this population, we developed a landscape suitability model and range expansion estimates (2023 2033). We assessed landscape suitability for breeding trumpeter swans with a use-available modeling design based on GPS collar data (20191iJ2023) and derived home range information from swans in the western Great Lakes Region of the U.S. and Canada. We related occurrence and pseudo-absences to landscape-scale summaries of upland and wetland conditions in a logistic mixed-effects model. We spatially applied the model and incorporated these estimates into a range-expansion model that identified the likelihood of a 50-km grid cell being occupied by 2033. We summarized time-series citizen-science data within cells and related newly occupied range cells and unoccupied cells (2004 2023) to year, survey effort, the amount of suitable breeding landscapes, and distance to previously occupied cells in a logistic regression model. Landscape suitability was positively related to greater amounts of wetland perimeter and foraging areas. New range cells were positively associated with suitable breeding landscapes and survey intensity and negatively associated with year and distance to previously occupied cells. We estimated a 4.4% (95% CI: 2.0-6.9%) annual range expansion rate from 2023 to 2033, with expansion occurring in the Prairie Pothole Region of the Dakotas and the Boreal Shield and James Bay Lowlands of Canada. North Dakota and South Dakota allow tundra swan (*Cygnus columbianus*) hunting but not for trumpeter swans. Our models can be useful for targeting hunter education at a local scale to mitigate unintended take of the similar looking species, allowing for pioneering trumpeter swans to establish persistent populations. Furthermore, our range expansion model can help jurisdictions with no or low breeding trumpeter swan populations anticipate future swan distribution and abundance.

---

During European colonization of North America, trumpeter swans (*Cygnus buccinator*) were extirpated from most of their historical range because of overharvesting and habitat loss (Banko 1960). The historical breeding range of trumpeter swans is believed to have extended approximately from Alaska, USA southeast to Newfoundland, Canada, southwest to Ohio, USA and west to Oregon, USA (Monson 1956; Banko et al. 1960). Trumpeter swan historical wintering range included suitable open water areas within their breeding range and throughout the remainder of the United States (Monson 1956; Banko et al. 1960). By the 1900s, their distribution was mainly restricted to the Greater Yellowstone Ecosystem and Alaska, USA (Banko 1960; Hansen et al. 1971).

Widespread efforts to reintroduce trumpeter swans to their historical breeding ranges commenced in the western United States in the 1930s to reestablish Pacific Coast and Rocky Mountain populations and within interior regions of the United States and Canada in the 1950s to reestablish an Interior Population (IP; Banko 1960). From the 1980s to 2000s, states and provinces in the upper Midwest region of the United States and southern Canada contributed extensive effort towards restoring trumpeter swans to their historical breeding range, with Minnesota, USA having the longest reintroduction history (Shea et al. 2002). Reintroduction efforts have been highly successful, and in 2015 the North American population was estimated to be ∼63,000 trumpeter swans with an average population increase of 6.5%/yr (1968 2015; Groves 2017). Population growth has been reported throughout their restored breeding range from the North American Trumpeter Swan Survey, with the Pacific Coast Population increasing 5.5%/yr, the Rocky Mountain Population increasing 6.5% year, and the IP increasing 14.4%/year (1968 2015; Groves 2017).

The Mississippi and Central Flyway Council management plan for IP trumpeter swans had a recovery objective of at least 2,000 swans with 180 successful breeding pairs by 2001 in the Central and Mississippi Flyways (Ad hoc Drafting Committee for the Interior Population of Trumpeter Swans 1998). These management goals have been greatly exceeded. During 2000 2015 the annual population growth rate for the IP was 19.9% and by 2015 their population was estimated to have ∼27,000 swans. Based on surveys in 2015 (Groves 2017), the bulk of the IP occurred in Minnesota (*n =* 17,021), Wisconsin (*n =* 4,695), Michigan (*n =* 3,021), and Ontario (*n =* 1,471), and based on 5-year estimates from 2000 2015 their respective annual population growth rates were 25%, 31%, 16%, and 11% (Groves 2017). As the IP has increased, so has its range. For example, in Ontario from 1991 to 2010, trumpeter swan range increased by ∼4.5 million ha (Handrigan et al. 2016). Since the last report (i.e., Groves 2017), The North American Trumpeter Swan Survey has been suspended indefinitely (Vrtiska et al. 2024a) but continued individual state-based efforts indicated populations were still increasing. For example, Minnesota conducted a state-wide population assessment and by 2022 their population estimate was ∼52,000 swans (Herberg and Rettler 2022), which is equivalent to a 17% geometric annual population growth rate based on the previously reported 2015 estimates (Groves 2017). Wisconsin has also conducted state-wide annual assessments since 2015 from spring Waterfowl Breeding Population and Habitat Survey and estimated 14,781 individuals in 2024, which is equivalent to a 14% geometric annual population growth rate based on previously reported 2015 estimates (Groves 2017; T. Finger, Wisconsin Department of Natural Resources, unpublished data).

Recent reports indicated that breeding swans are starting to pioneer into neighboring states in the Prairie Pothole Region including eastern North Dakota and eastern South Dakota that did not have reintroduction programs (Fisher et al. 2017; R. J. Murano, South Dakota Game, Fish and Parks, personal communication). These pioneering birds are presumably from the westward expansion of the IP in the Mississippi Flyway, given high population growth rates in Minnesota and lower growth rates in the Central Flyways High Plains flock of Nebraska and western South Dakota, USA. Although natural range expansion within their historical range is certainly an intended and welcomed consequence of reintroduction efforts, it does necessitate communication and planning to prevent conflict. For example, both North and South Dakota, USA have a hunting season for tundra swans (*C. columbianus*) and there is not a hunting season for the similar looking trumpeter swan. As such, managers need to have a better understanding about the potential expansion of their range into these areas to assist in future planning efforts.

To aid conservation planning, our goals were to develop a landscape suitability model for breeding IP trumpeter swans and highlight areas of future range expansion. We developed landscape suitability models with GPS collar data from IP trumpeter swans (Wolfson 2024; Wolfson et al. 2024), and wetland and land-cover data derived from Sentinel-2 remote sensing imagery. We estimated future expansion with long-term structured (i.e., North American Breeding Bird Survey; Sauer et al. 2017) and semi-structured survey data (i.e., eBird and iNaturalist; Sullivan et al. 2009; http://www.inaturalist.org). Our models could help conservation planning such as proactive human-wildlife conflict mitigation (e.g., unintended take), identifying landscapes suitable for trumpeter swans that could be targeted for protection, or identify areas that could be targeted for additional reintroduction. This information will also be useful as the Central, Mississippi, and Atlantic Flyway Councils currently work to update the IP Trumpeter Swan Management Plan and for jurisdictions that may experience future colonization by breeding swans or increases in swan distribution.

## STUDY AREA

Our study area for developing a landscape suitability model and predicting future range expansion represented a bounding box of trumpeter swan locations derived from GPS collars (Fig. 1) expanded to include the remainder of North Dakota and South Dakota. This bounding box included the upper Midwest and southern Canada, extending from the Rainwater Basin of Nebraska, USA in the southwest (99.5°W 40.3°N), to the shores of the James Bay of Ontario, Canada in the northeast (80.7°W 53.9°N). We included the remainder of North Dakota and South Dakota to better understand potential suitable breeding landscapes and range expansion into these areas, which likely will support breeding trumpeter swans in the future. The study area encompasses portions of four major ecoregions (Fig. 1; CEC 1997). This study area has a high diversity of wetland sizes and hydrological regimes (i.e., water permanence classifications), including major rivers such as the Mississippi and Missouri rivers, major lakes, such as the Great Lakes and Lake Winnipeg, many smaller lakes and peat bogs in the Northern Forest region (i.e., Minnesota and Boreal Shield), many smaller depressional wetlands of various hydrological regimes in the Great Plains (i.e., Prairie Pothole Region), and a great amount of shallow emergent wetlands in the Hudson Plains (i.e., James Bay Lowlands; CEC 1997). Upland cover is mainly comprised of grassland, cropland, and development in the southwest (i.e., Great Plains) transitioning into forest in the northeast (i.e., Eastern Temperate and Northern forests; CEC 1997, Brown et al. 2022; Fig. 1). Climate in this region varies from continental in the south to subarctic in the north (Beck et al. 2018). Although this study area remained the focus for the range expansion analysis, we extended the bounding box when collecting survey data (e.g. eBird data) for the range expansion analysis. This provided a broader geographic context of range expansion relative to our study area, and included data that could represent potential immigration into the study area from nearby regions, with specific interest on potential linkages between the High Plains flock of the IP and the remainder of the IP. This flock represents a subpopulation of the IP in Nebraska and South Dakota thought to be isolated from the rest of the IP (Comeau and Vrtiska 2011, Wolfson 2024). We expanded the bounding box west to the Rocky Mountains in Colorado (-107.5°W), south to the U.S.-Mexico border in Arizona (31.8°N), and east to the coast of southern Maine (-70.8°W).

**Figure 1.**
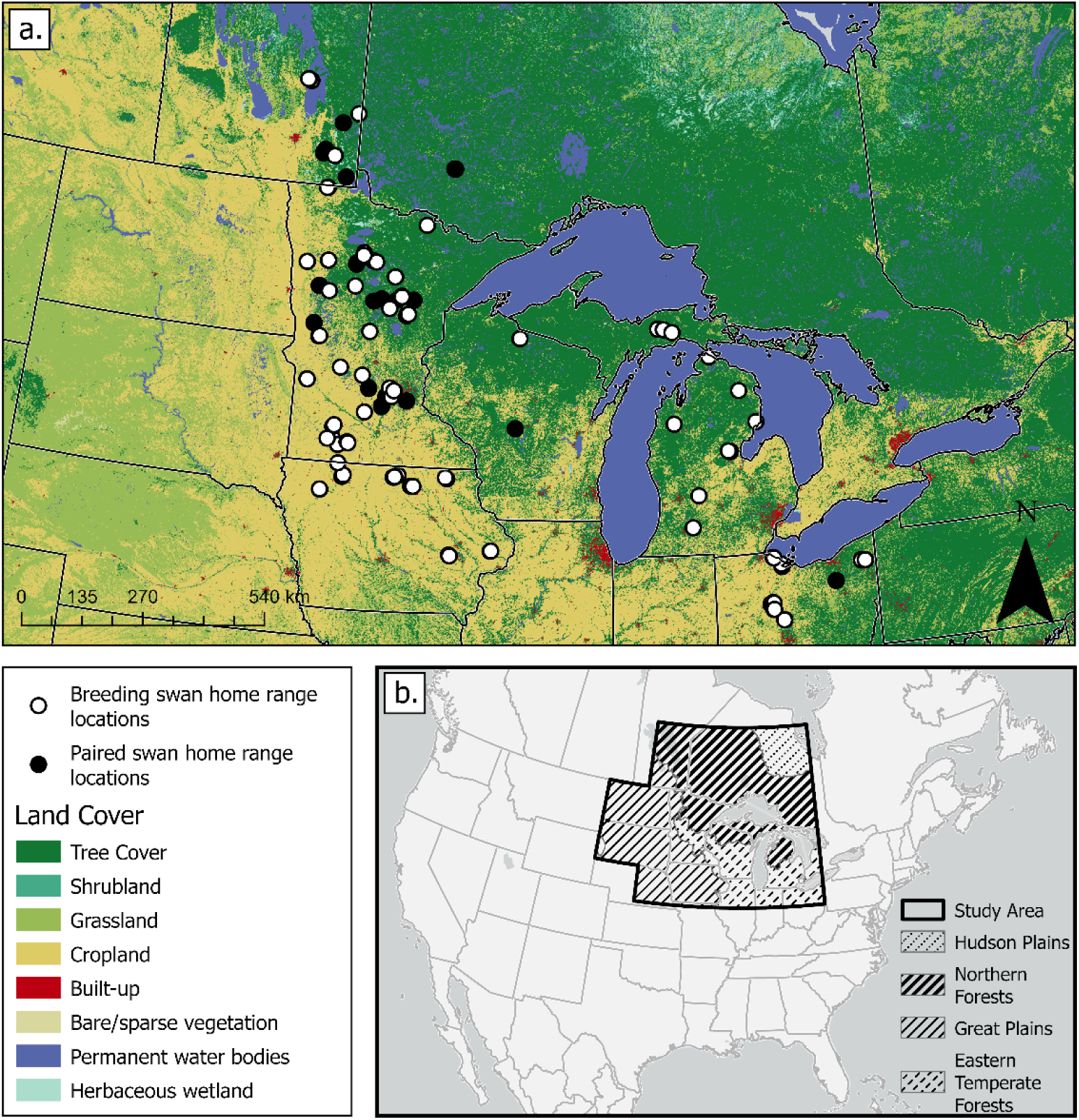
We estimated trumpeter swan home ranges from GPS-collar data (2019–2023) for breeding and paired swans within the western Great Lakes region of North America for each swan-year of data. Home-range data from swans that were known breeders were used in a home-range analysis and landscape-suitability model, whereas home-range data from paired swans where breeding status was unknown were only used in the home-range analysis. Home-range locations are shown in relation to land cover (panel a) and major ecoregions within our study area (panel b).

## METHODS

### GPS collar data

We developed a landscape suitability model with GPS collar data and information from utilization distributions (UDs) derived from these data. We obtained telemetry locations from 93 paired or breeding IP trumpeter swans (Table S1, Fig. S1 available in Supplemental Information). Wolfson et al. (2024) fit 1 swan per pair with GPS-GSM collar transmitters as part of a study of IP trumpeter swan movement ecology (Wolfson et al. 2024). They captured swans from 2019 through 2022 during the breeding season across 6 states and 1 province (Table S2, Fig. S1) and marked swans with 2 types of GPS transmitters incorporated into neck collars (see Wolfson et al. 2024 for details). We used GPS data collected from these marked swans from 15 April to 1 October 20191iJ2023; a period where breeding swans are on territories (Wolfson et al. 2024). We used a total of 2,009,014 GPS locations across 5 states and 2 provinces (Table S2).

### Home range analysis

We analyzed GPS collar data to derive home ranges and core areas, which we then used in model development. We calculated UDs from GPS telemetry data for each individual for each year (henceforth swan-year) in the computational environment R with the package amt (Signer et al. 2019, R Core Team 2024). We derived the 95^th^ isopleths from each UD to represent the individual’s home range and derived the 50^th^ isopleths to represent areas of higher use within the home range (i.e., core areas). We calculated speed, average sampling rate (i.e., time measurement between 2consecutive swan locations), and net squared displacement (i.e., distance from origin) for each swan-year. We removed tracks that had less frequent sampling rates (i.e., median > 1 hour) and thinned tracks with a resampling rate of 15 min and tolerance of 2 minutes (i.e., median and standard deviation of median values across tracks) to ensure tracks had similar sampling characteristics prior to developing home range estimates. We also removed locations with high speeds that could represent swans that were flying (i.e., >3 m/s), and removed points that represented relocation events that could occur from a failed breeding attempt (i.e., > 20 km from origin). The sample size of swan-year GPS locations varied due to when collars where deployed (e.g., during molt) and when they became inactive (e.g., failure). To ensure similar sampling durations for each swan-year prior to estimating UDs, we required that a minimum sampling duration be met. Additionally, to better understand the implications of temporal differences on space use, we also restricted sampling durations to 1of 3 periods during the breeding season. Ultimately, we developed up to 3 home range estimates per swan-year (hereafter breeding season period). These were estimated from ≥50 days of data prior to 30 June, ≥50 days of data after 30 June, and ≥25 days of data both before and after 30 June. These periods approximately align with ≥50 days of space use during the nesting period, or the brood-rearing and molt period, or both, respectively (Mitchell and Eichholz 2020). This approach allowed us to maximize the sample size of data for the landscape suitability model; for example, if we were unable to obtain a duration spanning nesting and brood-rearing/molt, we could use a home range from another period. Additionally, understanding how space use changes among these periods helped us understand the implications of using data from only a portion of the breeding period. Using the filtered GPS collar data, we calculated a UD with a continuous-time movement model and Ornstein-Uhlebeck foraging process (Fleming et al. 2014, Calabrese et al. 2016). This method represents random movement patterns with a tendency to return to a more central location (e.g., food source, nest).

Because space use could be influenced by individual and temporal differences that could then influence the landscape suitability model, we examined space use by sex, breeding status, and breeding season period. Breeding status included individuals that had reached sexual maturity (breeders) and individuals that were paired but sexual maturity was unknown. Paired individuals had a partner when captured but did not have cygnets or a known nesting attempt. Breeding individuals were paired and had cygnets when captured or had a known nesting attempt. Changes to breeding status in subsequent years were unknown, and an individual’s status remained fixed if they had multiple years of data. We used the R package glmmTMB (Brooks et al. 2017) to relate the size of home ranges and core areas to fixed effects for breeding season period, sex, and breeding status in a gamma log-link generalized linear mixed-effects model that included random intercepts for swan ID nested in year to account for subject-level and temporal variation. We then investigated marginal means and pairwise comparisons of factor levels with the R package emmeans (Lenth 2024). We computed estimated marginal means for factors based on the fitted model while accounting for all combinations of factor levels in the model and adjusting for multiple comparisons. To characterize how space use may shift over time, we calculated the distance between home ranges and core areas centroids across the 3 breeding season periods for each swan-year, respectively. Breeding season period comparisons included from early to middle periods, from middle to late periods, and from early to late periods. These relationships were characterized with the same modeling methods previously described. Lastly, we examined site fidelity for swans with more than 1 year of data by calculating the distance from core area centroids between all year-pairs (e.g. 2020-2021, 2021-2022, 2020-2022). For each swan-year we retained just 1 core area. If possible, we retained the home range and core area estimates derived from days spanning both nesting and molt/brood-rearing periods; if this was not possible, we used home ranges and core areas from the early breeding period (nesting), otherwise we used data from the late breeding period (molt/brood-reading). We report summary statistics for distance between core areas across all swan year-pairs.

### Trumpeter swan landscape suitability model: response variable

Home ranges and core areas guided the development of model training data under a use-available design (Johnson 1980, Manly et al. 2002) to develop a landscape suitability model (Guisan and Zimmermann 2000, Elith et al. 2006). We used these areas to filter occurrences, develop sampling areas for pseudo-absences, and inform the scales environmental data are summarized (i.e., window size for landscape-scale summaries of predictor variables). We only included data from individuals that were known breeders due to differences we found in space use between paired individuals with unknown breeding status and breeding individuals, and to better ensure we only characterized landscapes that are suitable breeding habitat. For each swan-year we retained data from 1 breeding season period using the same criteria implemented for the site fidelity analysis (i.e., prioritized selection as middle, early, then late period).

For each swan-year, we thinned the GPS collar data so all points were >100 m apart to reduce pseudo-replication (Steen et al. 2021). We retained all the thinned data points in the 50^th^ UD isopleth, and half the thinned data points outside the 50^th^ isopleth that were within the 95^th^ UD isopleth to ensure we characterized conditions in core areas but still had representation of conditions in areas of less use. For each swan-year, we generated a random sample of pseudo-absences that were equal in number to the occurrence points retained. These were placed randomly in an area represented by the home range buffered by 2,360 m (including the home range area), but not within 100 m of any known occurrence or other pseudo-absence. This buffer distance is 2 times the diameter of the median home range area (if represented by a circle) that we calculated across all individual home ranges used for developing model training data. This sampling strategy aligns with determining the local-and landscape-scale characteristics of areas used within a home range to other nearby available areas within and nearby the home range (Johnson 1980). The occurrence data used to restrict pseudo-absence sampling were based on all GPS collar locations (all swans and all years), and occurrence data obtained from eBird and iNaturalist citizen science databases, described below. Ultimately, we developed the model with data from 63 swans, representing 137 annual records (Table S1), with 7,934 occurrence points and 7,934 pseudo-absence points.

### Trumpeter swan landscape suitability model: predictor variables

We related occurrence and pseudo-absence data to 10-m land-cover data from the 2020 World Cover dataset (Zanaga et al. 2021) and the 20191iJ2023 Dynamic World dataset annually from 15 April to 1 October (Brown et al. 2022). These datasets are developed with Sentinel-2 remote-sensing imagery. World Cover is an annual global land cover dataset and Dynamic World is a land-cover classification dataset for each Sentinel-2 remote-sensing image. We processed these datasets in Google Earth Engine (Gorelick et al. 2017). We converted the 2020 World Cover dataset into 6 binary rasters where the value 1 represented a land cover of interest, otherwise 0. These land-cover types included grass, crop, development, forest, open water, and wetlands. We defined grass cover as a combination of grass, shrub, and bare/sparse vegetative cover classes (cover codes 10, 20, and 60). We processed the Dynamic World dataset by calculating the percentage of times a pixel was classified as flooded vegetation, and the percentage of times it was classified as open water. The revisit rate of a pixel is 2-5 days, with shorter intervals at greater latitudes due to swath overlap; however, this can be affected by factors such as cloud cover, which reduces revisit frequency by about half across latitudes (Brown et al. 2022). From these calculations we generated 4 binary rasters, each with the value 1 if certain conditions were met, otherwise 0. These represented areas that were classified as flooded vegetation with lower permanence (25-50% of the period) and higher permanence (>50% of the period), and open water with lower permanence (25-50% of the period) and higher permanence (>50% of the period). We combined these into 1 binary raster representing the full extent of the different water classes and then defined the wetland perimeter as a binary raster. We delineated perimeters with a zero-crossing edge detection technique (Marr and Hildreth 1980, Haralick 1984; Google 2023). Initially, we employed a difference-of-Gaussians filter to accentuate areas with rapid pixel value changes (i.e., change in sign), indicative of edges. This process converts the binary wetland raster into a continuous one. Subsequently, we identified wetland perimeters by identifying where the signs of pixel values transition from positive to negative or vice versa (Marr and Hildreth 1980, Haralick 1984, Google 2023). We thickened the perimeter using maximum value within a 3x3 pixel moving window to avoid data loss when projecting the raster (i.e., avoid perimeter values 1 being resampled to the value 0).

We extracted predictor variable values at occurrence and pseudo-absence point locations by buffering points by 3 radii and calculating proportional values of the binary rasters within buffer areas with the zonal statistics tool in ArcGIS Pro version 3.1 (ESRI, Redlands, CA). We used 50-m, 220-m, and 590-m radii. The first radius represents a local area of immediate surrounding. We selected the next 2 radii because circles with these radii approximately represented median core area and home range size, respectively, from the swan data used to develop training data. Ultimately there were 29 covariates for consideration in the landscape suitability model, which included all wetland-related covariates (*n* = 7) summarized at 3 scales, and all upland-related covariates (*n* = 4) summarized at the 2 larger landscape-scales. For all covariates retained in the final model after model selection, we calculated moving window statistics on their binary land-cover-class rasters at relevant landscape scales with the focal statistics tool in ArcGIS. For example, if the proportion of higher permanence flooded vegetation at 590-m radius landscape-scale was retained as a covariate in the final model, we conducted a focal statistics analysis on the higher permanence flooded vegetation binary raster with a 590-m radius circular moving window. We applied the model to the raster outputs from the focal statistics analysis to predict landscape suitability across the study area. Because Dynamic World variables were summarized by year and temporally matched to swan occurrence and pseudo-absence data, we spatially applied the model to covariate rasters developed with the same methods but based on data from 15 April to 1 October across all years (2019 2023) to represent average conditions during this period.

### Trumpeter swan landscape suitability model: model selection and validation

We modeled landscape suitability with a logistic mixed-effects model in R with the glmmTMB package (Brooks et al. 2017). We centered and scaled all covariates by subtracting the mean and dividing by the standard deviation. Random intercepts included swan ID nested in year to account for subject-level and temporal variation. We used an exploratory model-selection approach that implemented a constrained form of an all-model-subset method through the dredge function in the R package MuMIn (Bartoń 2023). We held random effects constant across all model subsets and fit models for all combinations of 5 uncorrelated fixed-effects covariates (i.e., Pearson’s |R-squared| < 0.7). We performed model selection using ΔAICc <2 to retain the most supported model (Burnham and Anderson 2003, Arnold 2010). We used the DHARMa package (Hartig 2022) to plot scaled residuals versus fitted values to assess uniformity. We reduced patterns in residuals by testing all possible 2-term interactions and quadratic terms for each covariate and evaluated performance with ΔAICc, marginal and conditional R-squared with the R package performance (Lüdecke et al. 2021), and reexamined scaled residual versus fitted values for improved fit. For the original top model from model selection and the updated model with reduced patterns in residuals, we present beta estimates and 95% confidence intervals for all covariates, marginal and conditional R-squared, and marginal effects plots that were derived with the R package sjPlot (Lüdecke 2024). We also estimated performance for these models with repeated 10-fold cross validation with 10 repeats. Strata for splitting data into training and testing folds were defined with occurrence and year combinations. We used observed and fitted values from population-level estimates (marginal estimates) to calculate area under the receiver operating characteristic curve (AUC) and Brier score for each repeat, and present mean AUC and Brier scores across the repeats. The AUC measures the model’s ability to discriminate between positive and negative outcomes, with higher values indicating better performance. Brier score is a measure of error calculated as the mean squared difference between predicted probabilities and observed outcomes. We spatially applied the model to generate population-level estimates (marginal estimates) across the study area.

### Range expansion model: response variable

We developed a logistic-regression, range-expansion model to make spatially explicit predictions of which geographical units across the study area will be part of the IP trumpeter swan’s breeding range by 2033. The model’s response variable was an annual binary indicator associated with 50-km hexagonal grid cells across the extent of the bounding box, which indicated if a cell was part of the swan’s breeding range (henceforth range cells). We conducted these analyses in R with the sf and tidyverse packages (Pebesma 2018, Wickham et al. 2019, Pebesma and Bivand 2023). We first determined range cells by summarizing time-series, citizen-science data within each cell, where cells were given the value 1 if they contained a trumpeter swan occurrence record, otherwise 0. With the runner package (Kałędkowski 2024), we then examined the time series, and when range cells reached 2 annual occurrences within a 6-year period, we assigned an indicator value of 1, otherwise 0. We did this to limit false positives that could be due to wandering individuals. Once we identified a range cell it remained a range cell in subsequent years.

We obtained trumpeter swan occurrence data from The North American Breeding Bird Survey (BBS; Sauer et al. 2017), eBird (Sullivan et al. 2009), and iNaturalist (http://www.inaturalist.org) databases. We obtained BBS route level data from 1966 2022, where route-level summaries are attributed to start locations. The eBird database contains records of bird observations (i.e., bird checklists) that are associated with point locations. The iNaturalist database has 1 record per observation that are most often photo documented and can have location uncertainty estimates.

We filtered these data in various ways prior to developing range models. We restricted all data to a bounding box larger than our study area (see Study Area section). For BBS data, we retained all surveys with a trumpeter swan observation along a standard route. For the eBird and iNaturalist dataset, we selected observations from June and July. For the eBird dataset, we removed any records that were deemed flyovers (behavior code) or from nocturnal flight calls (survey method). In addition, we read the comments of all records that had the word fly, and removed those that indicated the observation was a flyover. We did not use eBird records that had a travel distance > 10 km to better ensure observations were within the 50-km hex grid. For the iNaturalist data, we excluded records that had location uncertainty estimates >10 km. Ultimately, we used 231 observations in the BBS dataset (annual occurrence range: 1983 2019), 17,765 observations in the eBird dataset (annual occurrence range: 1957 2023), and 1,388 observations in the iNaturalist dataset (annual occurrence range: 1994 2023) to determine range cells over time. We used data from 2004 2023 in the range expansion analysis so that results are more aligned with natural expansion than the previous period of reintroductions.

### Range expansion model: predictor variables

We used 4 predictor variables in the range expansion model including year, the amount of suitable landscapes for breeding swans within each cell, the distance from each cell centroid to nearest range-cell centroid from the previous year, and log effort. We derived suitable landscapes from the landscape suitability model by converting the model to a binary output where values ≥ 0.84 were given the value 1, otherwise 0. We determined this threshold value with the cutpointR package (Thiele and Hirchfeld 2021), which examined observed and predicted values from 10-fold cross-validation of the landscape suitability model to determine a threshold value that limits false positives. We then summarized the proportion of suitable breeding landscapes in each grid cell. We included distance to nearest previously occupied range cell because we hypothesized cells closer to previously occupied cells could have a greater likelihood of being occupied in the future due to high adult and natal site fidelity (Lockman et al. 1985, Vrtiska et al. 2024b). We only included grid cells within our study area in the model, but the distance from newly occupied range cells to the closest previously occupied range cell could be calculated from range cells outside this area. Log effort was represented by the log-transformed number of eBird checklists submitted in the grid cell that year for the months of June and July, which we included to correct for the influence of effort being spatially biased. We summarized effort from the eBird sampling event dataset, which is a dataset of all checklist metadata but not their associated bird observations. We filtered the eBird checklists based on the same previous criteria (without filtering flyovers by comments or behavior code) and only kept 1 checklist if there were duplicated checklists (i.e., shared group checklists). We conducted these analyses in R with the terra (Hijmans 2024), sf, and tidyverse packages.

### Range expansion model: model validation, tuning, and forecasting

Prior to developing a spatially explicit range expansion model we first developed a simple model to estimate annual range expansion rate (area/year) across the study area that could be used to tune the output of the spatially explicit range expansion model (Hill et al. 2001, Preuss et al. 2014). With a gamma log-link generalized linear-regression model, we related the annual sum of range-cell area to year and the log transformation of mean annual effort across grid cells. We also tested a quadratic term for year because graphical plots indicated expansion may be decreasing in later years, and this may also be expected once suitable habitat becomes more limited. We also tested log effort as an offset, which scales the response variable. We included effort because some of the expansion is likely attributed to increased effort over time and not accounting for effort can bias estimates (Preuss et al. 2014, Zhang 2020). We used ΔAICc<2 to guide model selection, and assessed model fit with Nagelkerke’s pseudo-R^2^ (Nagelkerke 1991, Lüdecke et al. 2021). We calculated the geometric rate of range expansion from 2004 2023 and from 2023 2033 by making predictions for the start and end period, including 95% CI, and calculating the annual geometric growth rate (percent/year) for fitted values and lower and upper bounds. These values provided an understanding of past and potential future expansion, which we used to tune the output of the spatially explicit range expansion model.

We fit a logistic regression range expansion model to determine the likelihood that a grid cell is part of the IP trumpeter swan breeding range by 2033. For each year (2004 2023), we used newly occupied range cells (value 1) and an equal number of randomly selected non-range cells (value 0) as binary response variable in the model. There were 382 new range cells and 382 unoccupied cells in the model. We used AICc values to determine if a quadratic term for year improved model fit. We validated the model with observed and predicted values from 10-fold cross validation to calculate AUC and Brier score. We estimated IP trumpeter swan breeding range by 2033 by making annual predictions from 2023 2033 for non-range cells. For each year we converted the prediction to a binary output, where likelihood values > 0.5 were given the value 1, otherwise 0. We then accumulated range cells by merging each year’s newly predicted range cells to the previously occupied range cells and then updated distance values to previously occupied grid cells before making the next year’s prediction. We held the effort term constant across the study area when making predictions and ran the forecasting algorithm several times, testing a range of fixed effort values to determine the values that generated future range cells that were approximate to what would be expected from the previous range expansion rate analyses (i.e., area/year; fitted, upper bounds, and lower bounds).

## RESULTS

### GPS collar data and home range analysis

We used data from 93 swans collared by Wolfson et al. (2024), which generated 220 swan-years of data (Table S1). Prior to calculating UDs, we removed 62,165 GPS locations that were deemed flying or large relocation events and removed 2,198 locations when temporally resampling tracks. We removed 9 swan-years due to inadequate sampling intervals and removed 7 individuals and 37 swan-years due to inadequate sampling duration. Of the 86 swans for which we developed UDs, 59 had >1 year of data, resulting in 174 swan years of data (Tables S1 and S3). Of these 86 swans, there were 63 that were known breeders (137 swan years) and 23 that were paired with unknown breeding status (37 swan years; Table S1). Females were collared more often than males (∼1.7 females to males; Table S1) and most of the data were from Minnesota (∼50%; Table S2) and 2000 2022 (∼90%; Table S3). For approximately half of the swan-years we were able to develop UDs for all 3 periods (Table S4), and ∼ 90% had a UD developed for the late breeding season period (Table S4).

An analysis of pairwise comparisons (Table S5) of marginal means (Table S6) from models that related core area and home-range size to sex, breeding status, and breeding season period (Table S7) indicated that the size of core areas in the early breeding season period were 0.56 (95% CI:0.39–0.82; P = <0.001; Table S5) times the size of late breeding season period core areas (x [95% CI]: 84 ha [54–131 ha]; Table S6). Breeding swan core areas were 0.39 (95% CI: 0.17–0.91; P = 0.03; Table S5) times the size of paired swan core areas (x [95% CI]: 101 ha [48–214 ha]; Table S6), and breeding swan home ranges were 0.37 (0.17–0.84; P = 0.02; Table S5) times the size of paired swan home ranges (x [95% CI]: 658 ha [334–1406 ha]; Table S6). There was greater variation in estimates and uncertainty in differences between core area and home-range size by sex and by breeding season period (Tables S5 and S7).

An analysis of pairwise comparisons (Table S8) of marginal means (Table S9) from models that related the distance between core area and home-range centroids across annual breeding season periods to sex, breeding status, and breeding season period (Table S10) indicated the distance between early and late breeding season period core areas was 1.88 (95% CI: 1.60–2.21; P = <0.001; Table S8) times the distance between the early and middle breeding season period core areas (x [95% CI]: 189 m [123 – 283 m]; Table S9), and 1.98 (95% CI: 1.68 – 2.34; P= <0.001; Table S8) times the distance between middle to late breeding season period core areas (x [95% CI]: 177 m [116 – 268]; Table S9). Similar trends were found for distances between home-range centroids across annual breeding season periods (Tables S8 and S9). There was greater variation in estimates and uncertainty in differences between the distances of core-area or home-range centroids between early and middle breeding season period and middle and late breeding season period, and by sex or breeding status (Tables S8 and S10). An analysis of site fidelity between years indicated that the median distance between core area centroids of returning swans (i.e., >1 year of data) was 330 m (10th–90th quantile range: 62.3– 2,595 m). The minimum and maximum distances were 5–27,877 m.

Given the number of large outliers for home-range and core-area size (Table S11), we used median area estimates to inform the size of sampling strata for pseudo-absences and landscape-scale summaries of environmental conditions. The median home-range and core-area sizes used for model development, derived from breeding trumpeter swans for either the early, middle, or late breeding season periods, was 107.7 ha and 15.3 ha, respectively. This was similar to the median home-range (118.5 ha) and core-area (16.01 ha) sizes across all individuals (paired and breeding) and breeding season periods (Table S11).

### Landscape suitability model

We developed landscape-suitability models with data from 63 swans that were known breeders, resulting in 137 records (84 female and 53 male records; Table S1), with 43% of home ranges occurring in Minnesota (Table S2), and 89% of the data from 20001iJ2022 (Table S3). Approximately 58% of home ranges represented the middle breeding season period (*n =* 80), 35% represented the late breeding season period (*n =* 48), and the rest represented the early breeding season period (Table S4). There was 1 top model with a ΔAICc <2.The top model contained the variables proportion high-permanence open water and proportion low-permanence open water within a local area (50-m radius buffer), proportion grass within a smaller landscape-scale (220-m radius buffer), and proportion wetland perimeter and high permanence flooded vegetation within a larger landscape-scale (590-m radius buffer; Tables S12 and S13). All coefficient estimates were positive with 95% confidence intervals that did not include zero (Table S13). The greatest effect sizes were proportion high-permanence flooded vegetation and wetland perimeter (Table S13, Fig. S2). The model fit the data moderately well (marginal R2: 0.57; conditional R2: 0.72; Table S9) and had moderate performance across repeated 10-fold cross validation (x AUC: 0.83; x Brier score: 0.17). Model diagnostics indicated residuals versus fitted values had minor deviations from a normal distribution, with higher ranked residuals at moderate predicted values and lower rank residuals at higher predicted values. Both an interaction between flooded vegetation and perimeter and quadratic term for perimeter produced the greatest reduction in AICc values (Table S14). We retained both the quadratic and interaction terms in the final model, which resulted in an AICc that was 779.3 lower than the original model and produced a slight increase in fit and performance (marginal R2: 0.63; conditional R2: 0.77; x AUC: 0.84; x Brier score: 0.16; Table S14) and helped increase uniformity in residual versus fitted plots. The updated model had similar coefficient estimates as the previous model, but with higher predictions at mid-range values of perimeter when flooded vegetation was low and a weak positive linear association when flooded vegetation was high (Table 1, Fig. 2). All coefficient estimates had 95% CI that did not include zero (Table 1). The spatially applied model identified landscapes that had greater amounts of shoreline and flooded vegetation (Fig. 3). Spatial predictions of landscape suitability were high in areas of known occurrence but were also high across large geographies that did not have GPS occurrence records (e.g., Drift Prairie North Dakota, USA and James Bay Lowlands, Ontario, Canada; Fig. 3).

**Figure 2.**
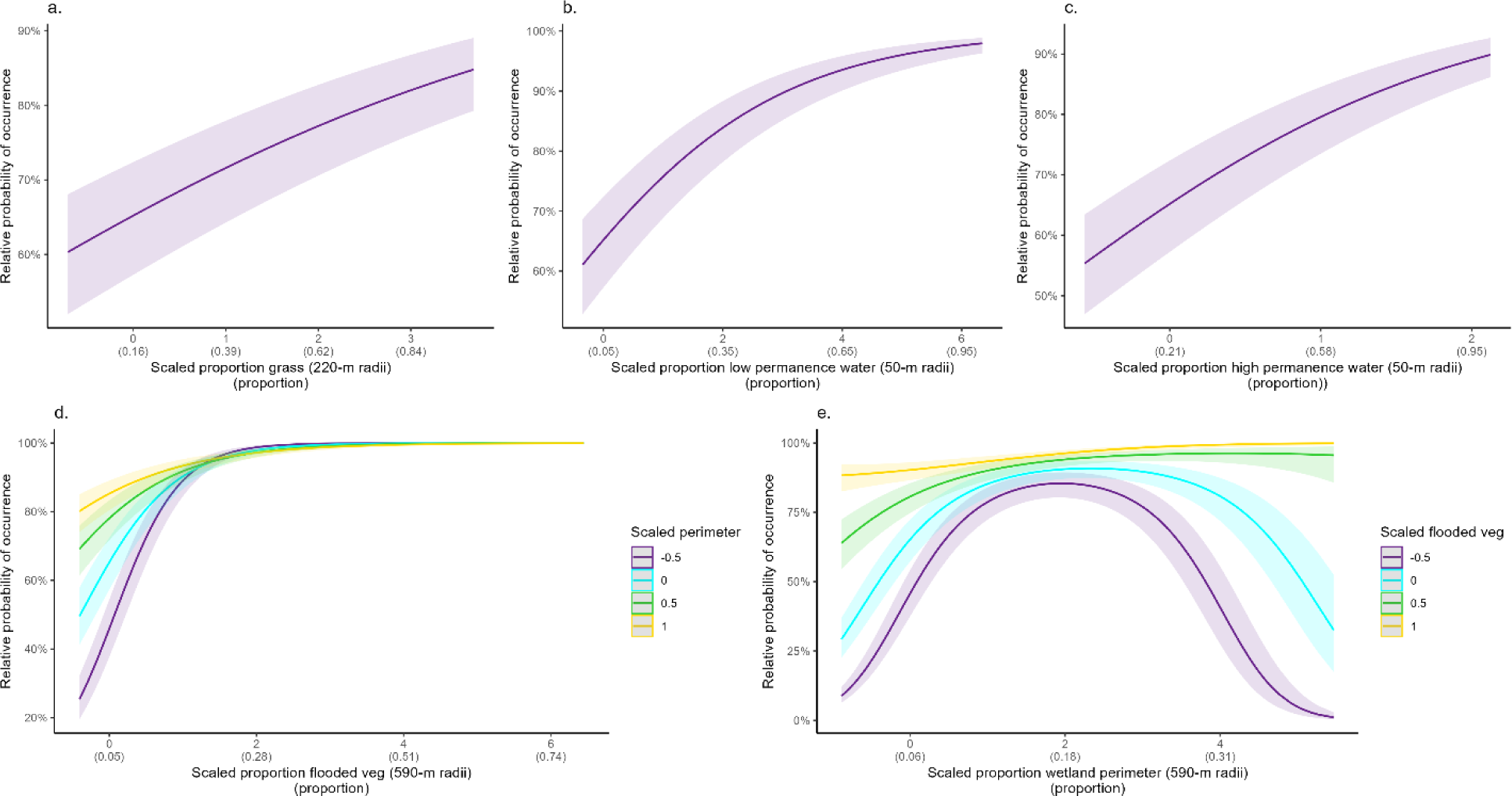
Marginal effects plots from a trumpeter swan landscape-suitability model showing relative probability of occurrence versus the following covariates: proportion grass (a), low-permanence open water (b), high-permanence open water (c), high-permanence flooded vegetation (d), and wetland perimeter (e). Shaded areas are 95% confidence intervals. The x-axis value labels include the scaled value (standard deviations from the mean) and their respective landscape-scale proportions. Variables with interactions have marginal effects plotted with the interaction covariate held at 4 different fixed values.

**Figure 3.**
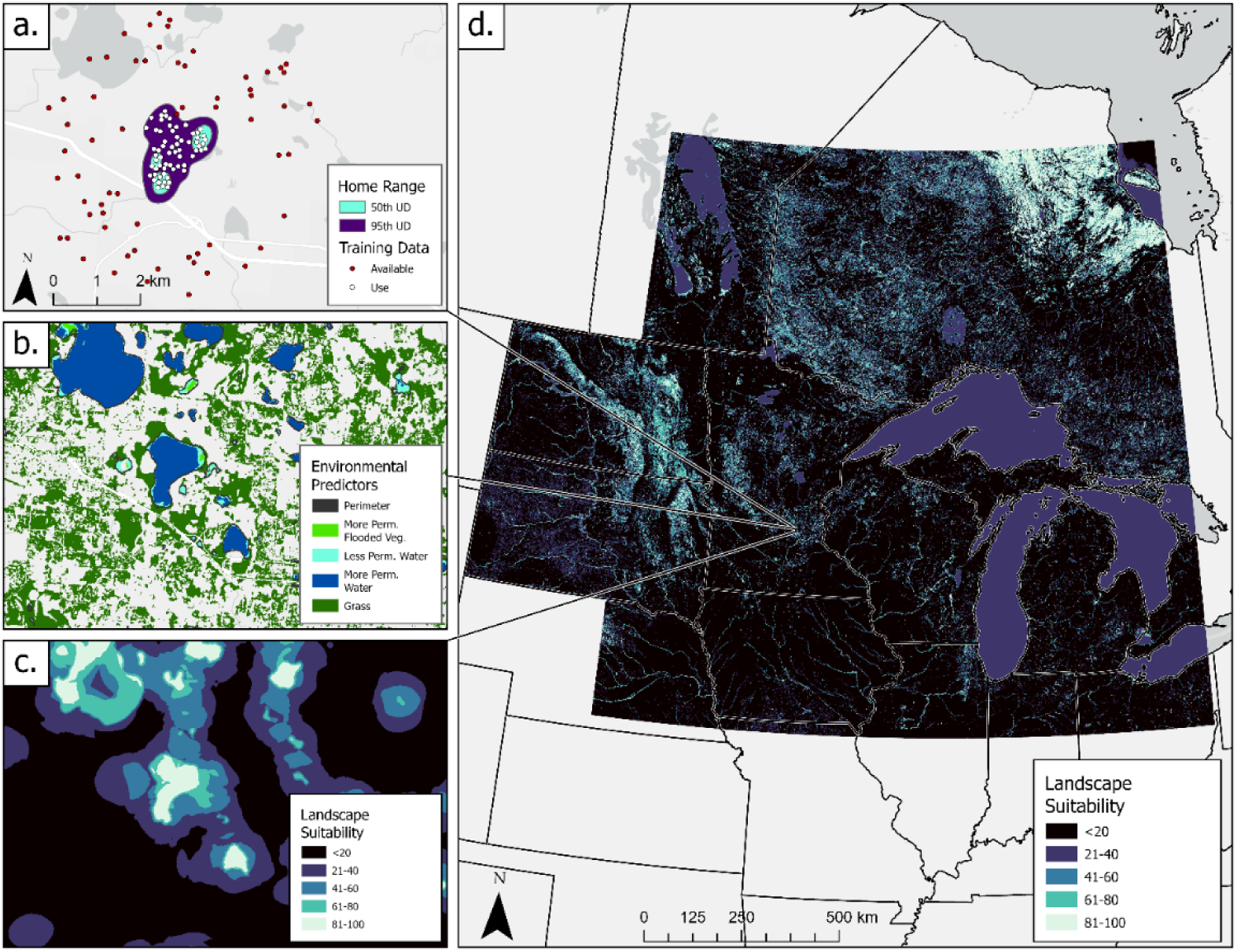
Trumpeter swan landscape-suitability model training data and spatial predictions for the western Great Lakes region of North America including home-range and core-area boundaries used to filter GPS collar data locations and define strata for pseudo-absence points (panel a), wetland and upland cover data that we summarized at landscape scales and used as covariate data (panel b), and model predictions (panel c) across the study area (panel d).

**Table 1.** Trumpeter swan landscape-suitability model summary. We developed the model by relating trumpeter swan occurrence and pseudo-absence data to local and landscape-scale summaries of the proportion of wetland and upland cover. We used an exploratory multi-model comparison for model selection and explored interactions and quadratics in the top model to reduce residual versus fitted patterns.

| Term | $\beta$ (95% CI) | SE | Z | P |
| --- | --- | --- | --- | --- |
| Intercept | 0.63 (0.29 – 0.96) | 0.17 | 3.67 | 0.002 |
| High permanence open water (50-m) | 0.73 (0.68 – 0.79) | 0.03 | 26.45 | <0.001 |
| Grass (220-m) | 0.30 (0.25 – 0.35) | 0.03 | 11.52 | <0.001 |
| Low permanence open water (50-m) | 0.51 (0.43 – 0.59) | 0.04 | 12.64 | <0.001 |
| High permanence flooded vegetation<br>(590-m) | 1.60 (1.48 – 1.72) | 0.06 | 25.79 | <0.001 |
| Wetland perimeter (590-m) | 1.43 (1.34 – 1.52) | 0.05 | 31.34 | <0.001 |
| Wetland perimeter<sup>2</sup> (590-m) | -0.31 (-0.34 – -0.27) | 0.02 | -16.89 | <0.001 |
| High permanence flooded vegetation<br>(590-m): Wetland perimeter (590-m) | -1.11 (-1.29 – -0.96) | 0.08 | -14.08 | <0.001 |
| High permanence flooded vegetation<br>(590-m): Wetland perimeter<sup>2</sup> (590-m) | 0.41 (0.33 – 0.48) | 0.04 | 10.68 | <0.001 |

### Home range expansion

The number of range cells and survey effort increased 2004 2023 (Figs. 4, 5, and S3). The top gamma log-link generalized linear-regression model included a quadratic term for year and log mean effort. Range area had a positive association with effort and time, with a weaker effect in later years (Table S15, Fig. 5). The model fit the data well and had a Nagelkerke’s R^2^ of 0.99. Modeled predictions 2004 2023 indicated range area increased from 85,084 km^2^ (95% CI: 81,607–88,709) in 2004 to 925,464 km^2^ (95% CI: 888,681–963,769 km^2^) in 2023 (Fig. 5), equivalent to a 13.38% (95% CI: 13.38% – 13.39%) annual increase and a total change in area by a factor of 10.88. When extrapolating predictions from 2023 2033 at the mean effort level in 2023, the range increased to 1,426,348 km^2^ (95% CI: 1,083,537–1,877,618 km^2^) in 2033 (Fig. 5), equivalent to an annual increase of 4.42% (95% CI: 2.00% – 6.90%) and a total change in area by a factor of 1.54. When effort was not included in the model the annual increase during 2023 2033 was 8.48% (95% CI: 6.74%–10.24%). When effort was included as an offset term, the model performed poorly (ΔAICc 52.88 from top model) and range area had a negative association with time.

**Figure 4.**
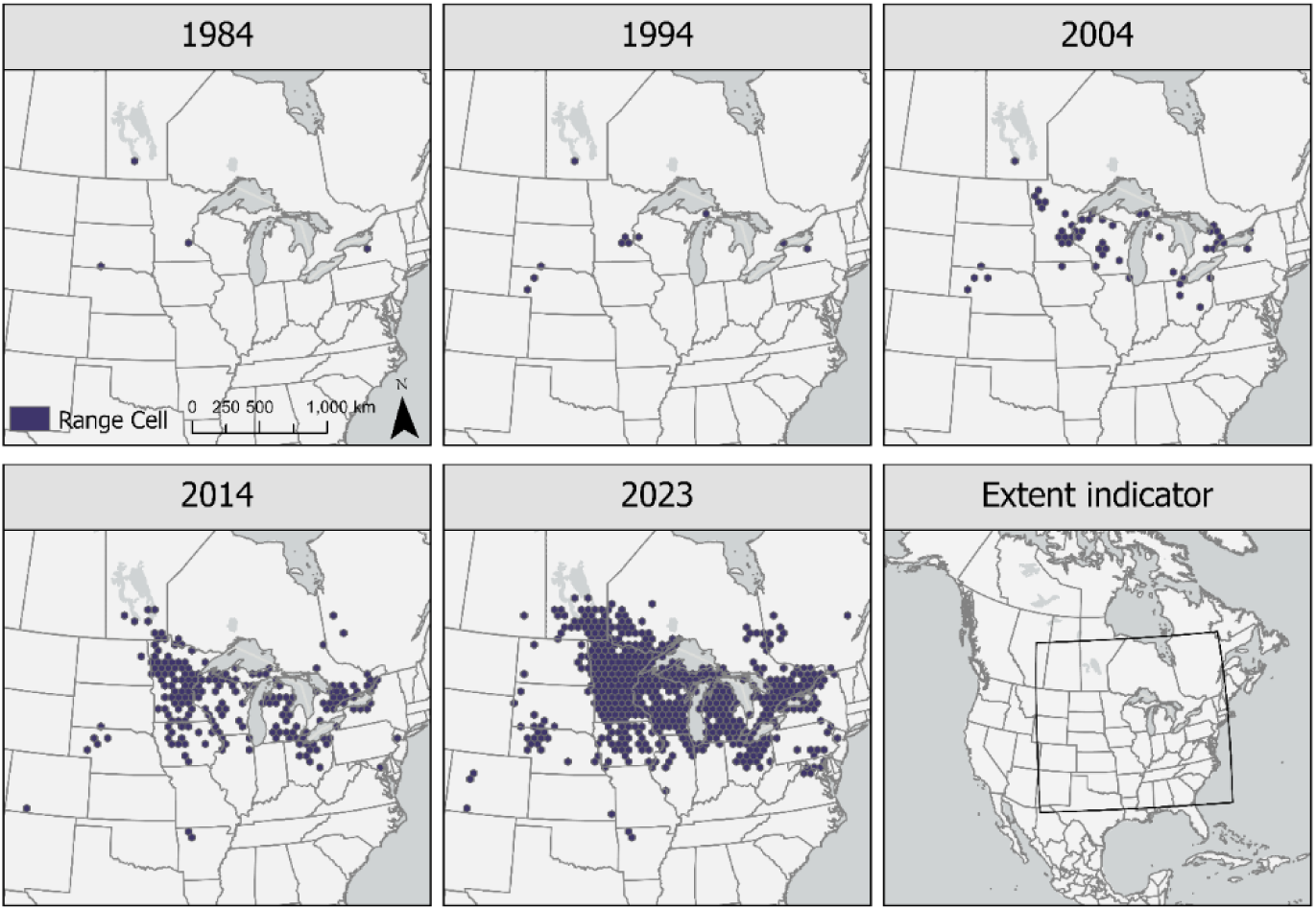
Trumpeter swan breeding range (1984–2023) depicted over 5 epochs in eastern North America. Range is represented as geographical units (50-km purple hex grids) that had ≥2 annual occurrence records within a 6-year period, which remained occupied into the future once reaching this threshold. We obtained occurrence data from long-term structured (i.e., The North American Breeding Bird Survey) and semi-structured citizen-science datasets (i.e., eBird and iNaturalist).

**Figure 5.**
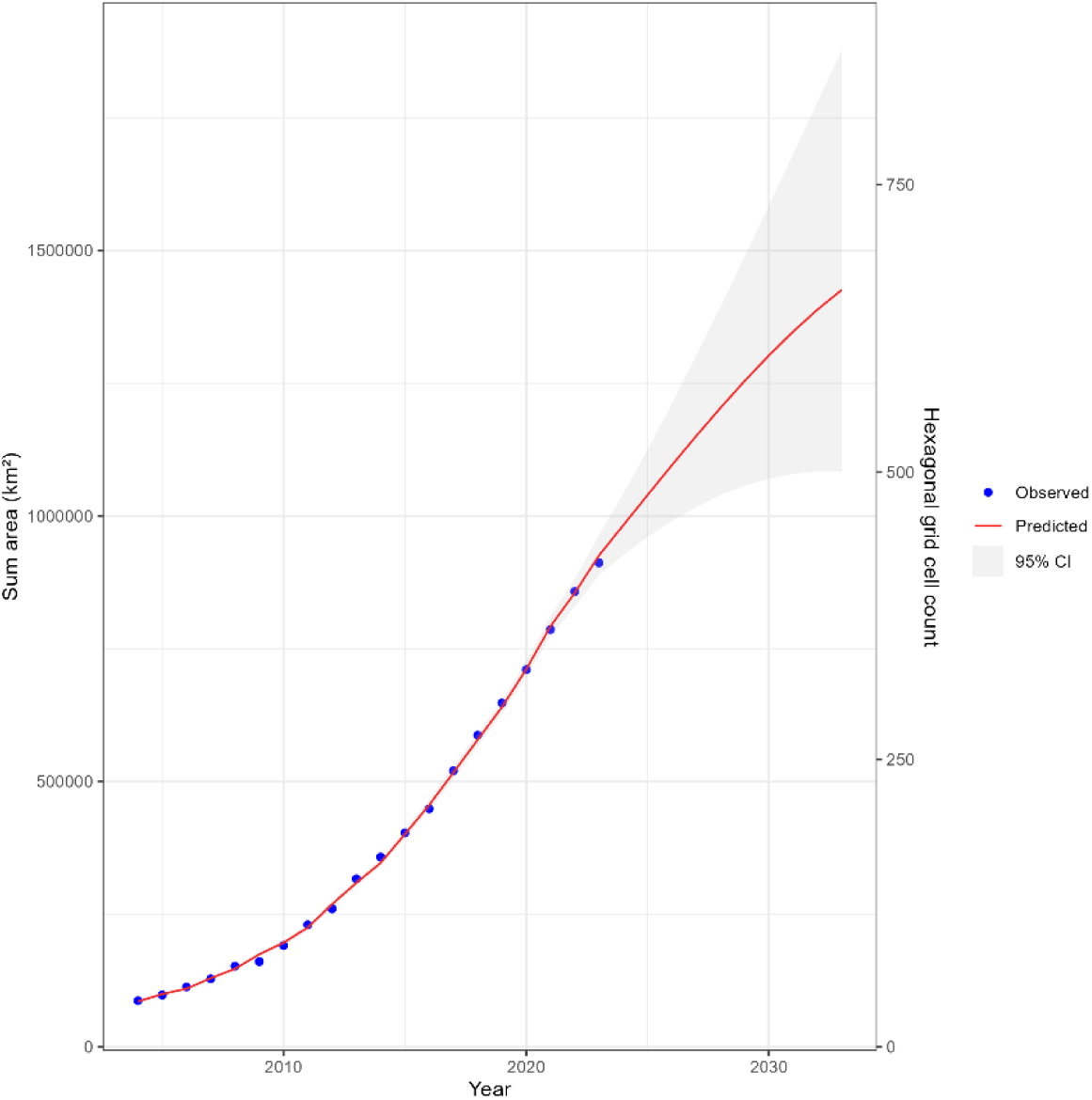
Observed and fitted values (2004–2023) and extrapolated future predictions (2024– 2033) from a range-expansion model that relates trumpeter swan range area (sum of geographical units occupied each year) in the western Great Lakes region of North America to year and survey intensity. We considered geographical units (50-km hex grids) part of the trumpeter swan range once it had ≥2 annual occurrence records within a 6-year period, and these units remained part of their range into the future once reaching this threshold. We obtained occurrence data from long-term structured (i.e., The North American Breeding Bird Survey) and semi-structured citizen science datasets (i.e., eBird and iNaturalist).

A quadratic term for year did not improve the logistic-regression model that made spatially explicit predictions of range expansion (ΔAICc=2). The model had a positive association with the proportion of suitable breeding landscapes (7.88 [95% CI: -0.45 – 14.87]), and log effort (0.53 [95% CI: 0.40 – 0.67]), and negative associations with year (-0.12 [95% CI: -0.16 – -0.07]) and distance to nearest previously occupied range cell (-0.02 [95% CI: -0.03 – -0.02]; Table S16). It had an AUC of 0.87 and Brier score of 0.14 calculated from 10-fold cross validation. At the mean estimated annual growth rate, range expansion was predicted to occur in the Boreal Shield and James Bay Lowlands of Canada and portions of the Prairie Pothole Region in the Dakotas, with less extensive expansion in Ontario and southwestern South Dakota (Fig. 6). At the lower bounding growth rate, range expansion was constrained to the portions of the Boreal Shield and a portion of the Prairie Pothole Region in the Dakotas, and less extensive expansion in Ontario (Fig. 6). At the upper bounding growth rate, range expansion was predicted to occur in the James Bay Lowlands and the 2023 range expanded by a ∼1-4 cell buffer throughout the study area (Fig. 6).

**Figure 6.**
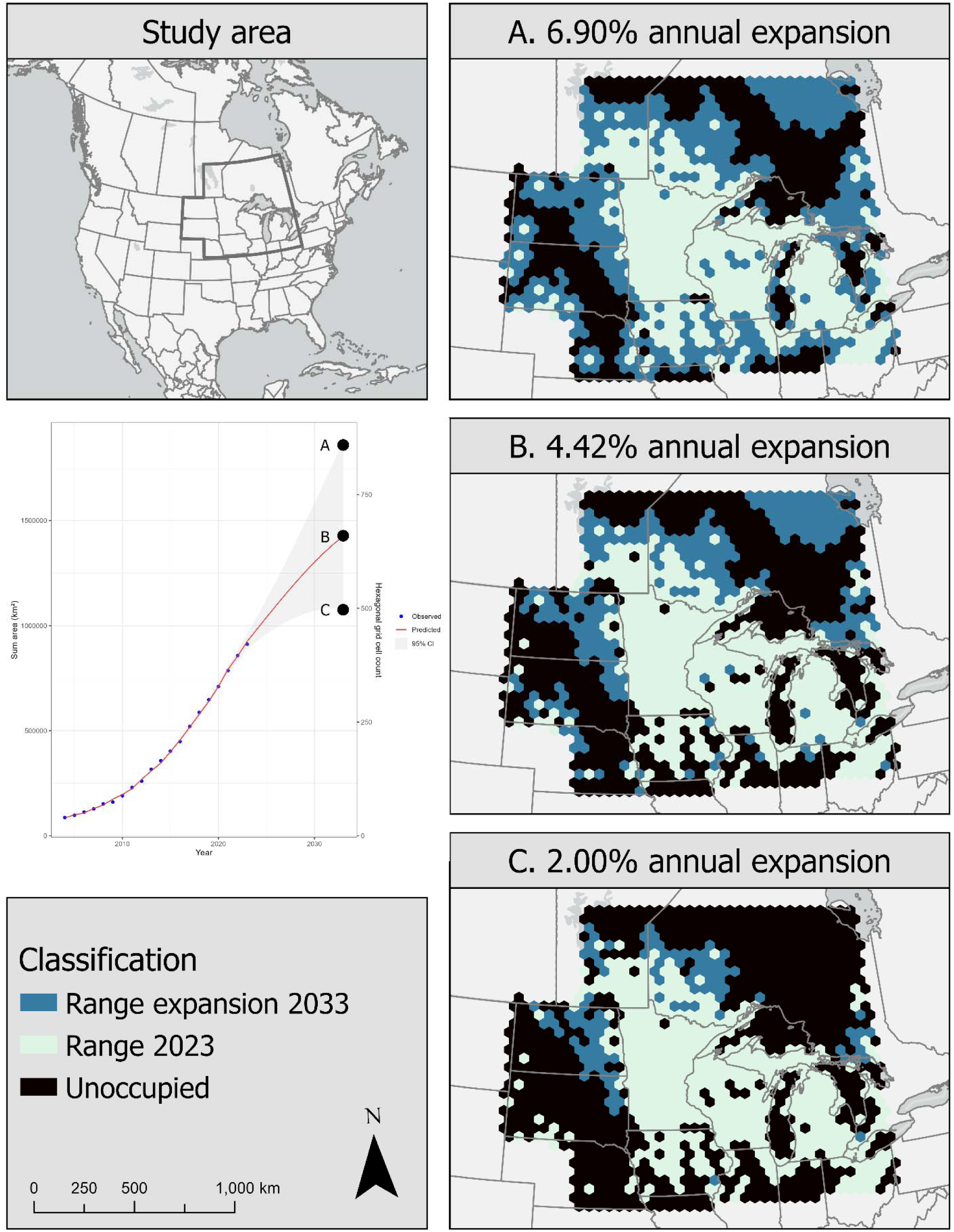
Trumpeter swan range in 2023 and predicted range expansion by 2033 within the western Great Lakes region of North America derived from a spatially explicit range-expansion model that relates newly occupied units and unoccupied units to suitable conditions within those units, distance to previously occupied units, year, and survey effort. When predicting into the future, survey effort was held at values that produced range-area estimates that align with predictions in 2033 (fitted lower and upper bounding 95% CI predictions) from a model that related range area to year (e.g., line plot and corresponding panels a, b, and c). We considered geographical units (50-km hex grids) part of the trumpeter swan range once they had ≥2 annual occurrence records within a 6-year period, and these units remained part of their range into the future once reaching this threshold. We obtained occurrence data from long-term structured (i.e., The North American Breeding Bird Survey) and semi-structured citizen science datasets (i.e., eBird and iNaturalist).

## DISCUSSION

Using GPS location and land-cover classification data, we characterized IP trumpeter swan space use and assessed landscape suitability across an extensive geographic area. Combining these data with an analysis of standardized bird survey data and citizen science observations, we estimated range expansion rates and identified regions where trumpeter swans are most likely to expand their breeding distribution over the next decade. These predictive insights can equip wildlife managers with the necessary information to anticipate trumpeter swan range expansion and devise strategies to mitigate potential wildlife management issues.

The environmental associations characterized in our landscape suitability model aligned with biological expectations. Trumpeter swans have been associated with open basins, such as lakes and ponds, that have complex shorelines and submerged aquatic vegetation (Banko 1960, Hansen et al. 1971). Our models indicated that landscape suitability increased with greater amounts of high-permanence flooded vegetation, grass, low-permanence open water, and wetland perimeter (Table 1, Figs. 2 and 3), which could represent complex shoreline regions that provide greater amounts of emergent or submergent vegetation foraged on by swans (Henson and Cooper 1993, Grant et al. 1994, Squires and Anderson 1995, LaMontagne et al. 2011) due to shallower water and increased light penetration. Additionally, a positive association with high-permanence open water could have represented areas with deeper water away from shore that would allow swans to safely retreat from terrestrial predators and other disturbances (Henson and Grant 1991).

Our models did not include climate variables because we wanted to avoid restricting predictions to an inaccurate or perceived climate envelope, which could result from the species not yet fully reoccupying its former range. Although it may seem reasonable to view predictions of highly suitable landscapes in currently unoccupied regions as over-predictions outside of their climate envelope, there is evidence that swans historically occupied these locations (e.g., Prairie Pothole Region of North Dakota, USA and James Bay lowlands of Ontario, Canada; Banko 1960; Houston and Houston 1997; Lumsden 1984). A status quo bias may temper the expectation that trumpeter swans will exhibit extensive range expansion; however, the reintroduction of another Anatidae, the giant Canada goose (*Branta canadensis maxima*), is an impressive example of the potential capacity for recovery after reintroduction efforts. This non-migratory subspecies of Canada goose was thought to be extinct until a small population was found in Minnesota in the 1960s (USDI 1963, Hanson 1965). Following reintroduction efforts (Nelson and Oetting 1998), their populations rebounded and resident, giant Canada geese now outnumber migratory geese to the point they are considered a nuisance species in some regions (Conover and Chasko 1985, USDI 2006, Dolbeer et al. 2014).

The data we used in the landscape suitability model were more representative of space use of breeding female swans in Minnesota later in the breeding season due to a greater number of collared females than males (Table S1), greater collared swan occurrence in Minnesota compared to other locations (Table S2), and 35% of the records representing home range estimates and GPS data from only the late breeding season period (Table S4). Our analyses did not find any sex-based differences in space use suggesting that having more collared females than males did not bias home-range or core-area estimates (Tables S5, S6, S8, and S9). Minnesota’s landscape, rich in lakes and diverse landcover types, is not representative of conditions throughout the modeling region (Fig. 1). A more geographically or environmentally representative training dataset could improve model predictions and enhance generalization or transferability of the model across varied landscapes and conditions (Charney et al. 2021). However, the rapid expansion of trumpeter swans in Minnesota may also reflect highly suitable conditions in this region, which has been characterized by the model. Core areas were smaller in the early breeding season, which could be associated with more restrictive space use because of territory establishment, mate guarding, nest building, and shared incubation duties (Henson and Cooper 1992, Lumsden 2018), whereas in the late breeding season period there may be increased stochasticity of space use associated with parental care of young and changing conditions or resource use and availability (Mitchell and Eichholz 2020). For potentially similar reasons, space use also shifted over time. Consequently, important nesting conditions during the early breeding season period may be under-represented in our model.

Because trumpeter swans can pair early but not breed until several years later (Banko 1960), our decision to exclude paired individuals with unknown breeding status from the model likely increased the model’s ability to identify suitable landscapes for breeding swans. However, it is unknown if any swans attempted nesting or successfully produced young after their initial capture and breeding status designation. It is thought that once trumpeter swans reach maturity, they will nest at 1-year intervals if conditions are suitable, but this has not been well studied for trumpeter swans; however, the probability of nesting in subsequent years for mute swans (*Cygnus olor*) has been documented at 90% (Birkhead and Perrins 1985, Ellis and Elphick 2007). Despite this uncertainty, paired individuals had larger core areas and home ranges, which could be associated with trying to find more suitable landscapes or perhaps not being as selective of landscapes where resources necessary for breeding are in proximity. This trend has also been reported for tundra swans in Alaska (Spindler and Hall 1991).

The goals of our assessment of trumpeter swan range expansion, the methodology we chose, and the characteristics of citizen science data presented several considerations. Many of our methodological choices aligned with providing results at spatial and temporal scales that are useful for conservation decision-making. The range expansion model is at a regional spatial scale and can be used to help understand the potential for trumpeter swan range expansion. The landscape-suitability model is at a smaller, local spatial scale, which can be useful for targeting wetland complexes or communities for conservation actions (e.g., protection, reintroduction, hunter education). Although we used GPS collar data to inform the appropriate spatial scales for the landscape-suitability model, there was not an optimal data-driven method for selecting the scale at which the extent of occurrence should be represented (i.e., grid size) in the range-expansion model. Given the size of the study area, a 50-km grid cell was visually appealing from a data presentation standpoint. It also provided some granularity for regional assessments within a state or province, and the grid-cell approach enabled discontinuity of range estimates that can help reduce commission and better support conservation decisions. This was also appropriate for a highly mobile species that is not restricted to a front-line type of range expansion. The criteria we implemented for establishing a range cell (2/6 years) also provided greater confidence that the grid cell was part of the range and reduced the likelihood a cell was considered occupied by wandering individuals, which is possible for a highly mobile species, and a data source that may have occurrence records represented by species flying overhead (easily detected and identified by their flight call). This decision limited the extent of occurrences and there were several cells that had 1 annual occurrence in a 6-year period that were often in areas of predicted range expansion, suggesting that our projections may underestimate range expansion.

Developing range-expansion models with citizen-science data had associated challenges. Citizen-science data are spatially and temporally biased; however, their sampling intensity, geographic coverage, and the competitive framework of these databases make them especially appealing for range-expansion analyses. Modeling efforts that use citizen-science data can improve their predictive performance by including effort terms that can account for biases. For example, in species-distribution models that are developed with eBird data, complete checklists (i.e., all species are reported) are used to develop presence and absence data and survey duration and travel distance can be used to account for varying effort. However, to capture the full extent of occurrences across a region, we did not limit our data to only include complete checklists and instead used complete checklists and incidental and incomplete sightings, which lack associated effort data. We therefore incorporated sampling effort (i.e., the number of checklists submitted in an area) as a covariate representing the amount of data available to determine an occurrence. Another option could be to directly account for detection probability in the models (Kéry et al. 2013) given repeated visits to a location (e.g., eBird hotspots). Although we did not estimate detection probability, our study benefited from the subject being a conspicuous bird that is large, loud, white, and often out in the open (flying in the air or floating on a waterbody). In addition, it was a regional rarity during its reintroduction, which may have increased the likelihood of documentation in competitive birding databases like eBird.

Trumpeter swan breeding range and eBird use were spatially and temporally correlated (Fig.6 and Fig. S4). Accounting for this correlation in the model improved predictive performance and decreased the predicted range expansion by approximately half. Surprisingly, incorporating effort as an offset in the model to scale the response variable resulted in estimates indicating a declining range area over time, which is counter to empirical observations. This discrepancy likely occurred because the effect of effort does not have a fixed proportional relationship with range area *vis-à-vis* detection of occurrence, potentially due to high detection and reporting of the species in citizen-science databases despite lower effort in early years. Effort had a significant effect on the spatially explicit range-expansion model. Specifying effort values that produced estimates in 2033 that were approximate to predicted future range area estimates seemed more reasonable than specifying the 2023 mean effort value in the model, which predicted full expansion across the entire study area over the next 10 years.

Although trumpeter swans are highly mobile, range expansion (i.e., new range cells) had a negative association with distance to previously occupied range cells, indicating that a front-line type of expansion may be occurring. This could be due to behavioral characteristics, such as natal site fidelity and shorter dispersal distances from natal sites, but to our knowledge there have been few studies on this topic for trumpeter swans. However, studies have reported wintering (Vrtiska et al. 2024b) and breeding site fidelity of adults and reported observations of family groups being maintained into winter and into the next breeding season (Lockman et al. 1985). Our results also support adult site fidelity in our study area. Incorporating landscape suitability in our model was helpful for identifying grid cells that are more likely to be occupied in the future. In some instances, this variable had a strong enough effect that expansion was predicted in areas at farther distances from previously occupied range cells (e.g., James Bay Lowlands). This frontier expansion would likely be contingent on how far swans wander to prospect new breeding locations and if environmental conditions not included in the landscape suitability model, such as ice-off or ice-on dates (i.e., season length; Schmidt et al. 2011), allow for populations to establish. The range expansion model also had a negative association with year that could indicate some level of saturation is already occurring.

### Conservation implications

The resurgence of any native species that was once widespread raises several ecological and management considerations. The potential ecological effects of a re-established and growing trumpeter swan population could include interspecific competition for local resources, particularly as the trumpeter swan breeding range expands, potentially overlapping the breeding range of tundra swans, which fill a similar ecological niche (Wilk 1993, Schmidt et al. 2011, Engelhardt et al. 2014, Kouzov et al. 2024). Additionally, trumpeter swan population increase could present challenges in management and human-wildlife conflict mitigation, similar to issues faced with Canada goose recovery (Conover and Chasko 1985). These challenges may include vegetation damage (Radtke and Dieter 2011, Washburn and Seamans 2012, Wood et al. 2014, Nilsson et al. 2016), aviation collisions (Dolbeer et al. 2014, Dolbeer 2020), the spread of zoonotic diseases (Gorham and Lee 2014, Elmberg et al. 2017), weed dispersal (Ayers et al. 2010), and degraded water quality due to increased nutrient loading (Manny et al. 1994, Post et al. 1998, Unckless and Makarewicz 2007).

Trumpeter swan range expansion could also have genetic implications for the High Plains flock by increasing genetic diversity and connectivity within and between reintroduced populations (Wolfson 2024). The High Plains flock was reintroduced in the 1960s and has remained isolated and exhibited low population growth and range expansion rates that have contributed to lower genetic diversity. If the larger IP breeding range expands to come in contact with the High Plains flock breeding range, there is the potential for increased gene flow and resulting greater genetic diversity and resilience or adaptability to environmental changes (Proft et al. 2018).

Waterfowl hunting considerations also play a potential role in the expansion dynamics of trumpeter swans, especially in the context of hunting tundra swans. With the expansion of trumpeter swans into jurisdictions that allow hunting of similar-looking tundra swans (i.e., North and South Dakota), the probability of incidental take of trumpeter swans will likely increase (Drewien et al. 1999, Fisher et al. 2017). In response to this increased likelihood and potential citation for illegal harvest, there has been a desire to limit the expansion of trumpeter swans into areas that have tundra swan hunting (Engelhardt et al. 2000, Lumsden and Drever 2001). To mitigate these issues, the U.S. Fish and Wildlife Service completed a General Swan Hunting Season Environmental Assessment in 2019 (U.S. Fish and Wildlife Service 2019) to establish a framework for hunting regulations to govern the take of both trumpeter and tundra swans in portions of the Atlantic, Mississippi, and Central Flyways within the range of IP trumpeter swans that currently have operational hunting seasons on Eastern Population tundra swans or may have in the future. To date, no states have implemented a general swan hunting season largely because they want to protect pioneering trumpeter swans from local extirpation (M. Syzmanksi, North Dakota Game and Fish Department, personal communication). Once trumpeter swan populations in some jurisdictions pass a certain threshold, however, it may be necessary to consider a general swan season as trumpeter swan populations continue to grow and the likelihood of unintended harvest increases. Alternatively, some jurisdictions may wish to emphasize hunter education efforts to minimize take of trumpeter swans.

## ACKNOWLEDGMENTS

The authors were funded by the U.S. Fish and Wildlife Service (KWB, TRC), the U.S. Geological Survey (DEA), and via a grant from the Minnesota Environmental and Natural Resources Trust Fund (DWW) and support from the Minnesota Cooperative Fish and Wildlife Research Unit (DWW and DEA). We thank the cooperators on the study of Wolfson et al. (2024) who participated in capturing and marking IP trumpeter swans: Michigan Department of Natural Resources; Ohio Department of Natural Resources-Division of Wildlife; Iowa Department of Natural Resources; the Canadian Wildlife Service Branch of Environment and Climate Change Canada; Wisconsin Department of Natural Resources; Great Lakes Indian Fish and Wildlife Commission; Minnesota Department of Natural Resources; Three Rivers Park District; Arkansas Game and Fish Commission; Manitoba Department of Economic Development, Investment, Trade, and Natural Resources; U.S. Department of Agriculture Animal and Plant Health Inspection Service; Trumpeter Swan Society; Cleveland Metroparks Zoo; Toledo Zoo; Cincinnati Zoo; and the American Association of Zoo Keepers chapters from the Akron, Columbus, Cincinnati, and Clevland areas. We thank D. Arsnoe, R. Batterson, L. Brown, S. Cordts, B. Davis, V. Drake, A. Duffiney, J. Fredrickson, M. Garrick, D. Hoffman, S. Hogg, J. Huener, J. Hummel, G. Ivey, D. McArthur, C. McCarty, L. Naylor, K. Norris, M. North, K. Rowe, R. Ruden, J. Schmidt, B. Shirkey, N. Smith, E. Thorson, G. Westerfield, S. Zaleski, and E. Zlonis for assistance with fieldwork. We thank B. Avers, F. Baldwin, K. Bekker, W. Brininger, P. David, W. Ford, D. Luukkonen, and H. Saloka for assistance with project logistics. We thank all who have conducted surveys and contributed observations to The North American Breeding Bird Survey, eBird, and iNaturalist databases and we thank staff that have helped develop and maintain these databases and made them publicly available. We thank H. M. Hagy, reviewers, and editors, whose comments greatly improved the manuscript. The findings and conclusions in this article are those of the authors and do not necessarily represent the views of the U.S. Fish and Wildlife Service. Any use of trade, product, or firm names is for descriptive purposes only and does not imply endorsement by the U.S. Government.

## ETHICS STATEMENT

Protocols for capturing and marking trumpeter swans in the United States were approved by the University of Minnesota Animal Care and Use Committee (protocol no. 1905-37072A), the U.S. Fish and Wildlife Service (Research & Monitoring Special Use Permits no. K-10-001, M-20-002, and M-21-014), the U.S. Geological Survey Bird Banding Laboratory (Federal Bird Banding Permit no. 21631) and state-specific permits approved by each state wildlife agency involved. All captures and marking of trumpeter swans in Manitoba were conducted under Federal Scientific Permit to Capture and Band Migratory Birds (no. 10271), Federal Animal Care Committee approval (project 20FB02), Provincial Species at Risk Permit (no. SAR20012), and Provincial Park Permit (no. PP-PHQ-20-016).

