## Supplemental Tables and Figures for "Landscape suitability and range expansion estimates for the North American Interior Population of trumpeter swans"

4/3/2025

Table S1. We developed home range (HR) estimates from trumpeter swan GPS collar data by individual for each breeding season that the collar transmitted data. These data were analyzed to better understand space use differences by sex and breeding status. Home ranges were filtered to develop landscape suitability models with swan data that were known breeders. We provide sample sizes for individuals and individual:year combinations, sample sizes after removing individuals with insufficient data for home range estimates (HR output), and sample sizes after removing home range estimates of paired individuals (HR filtered).

|  | Individuals | | | | |  | Individuals:Year | | | | |
| --- | --- | --- | --- | --- | --- | --- | --- | --- | --- | --- | --- |
|  |  | Sex | | Breeding Status | |  |  | Sex | | Breeding Status | |
| Group | n | Female | Male | Breeder | Paired |  | n | Female | Male | Breeder | Paired |
| Collar data | 93 | 59 | 34 | 70 | 23 |  | 220 | 132 | 88 | 175 | 45 |
| HR output | 86 | 55 | 31 | 63 | 23 |  | 174 | 109 | 65 | 137 | 37 |
| HR filtered | 63 | 39 | 24 | 63 | 0 |  | 137 | 84 | 53 | 137 | 0 |

Table S2. We developed home range (HR) estimates from trumpeter swan GPS collar data by individual for each breeding season that the collar transmitted data. Home range estimates were analyzed to better understand space use and were filtered to develop landscape suitability models with swan data that were known breeders. We provide sample sizes per state or province for the number of GPS locations, number of individuals captured and collared, number of home range estimates across individuals and years, and number of home range estimates from breeding swans whose data were used to develop landscape suitability models.

|  | State or province | | | | | | | |
| --- | --- | --- | --- | --- | --- | --- | --- | --- |
| Data | AR | IA | MB | MI | MN | OH | ON | WI |
| GPS locations | 0 | 288,636 | 149,024 | 263,937 | 954,587 | 286,175 | 48,856 | 17,810 |
| Individuals collared | 3 | 11 | 7 | 13 | 44 | 14 | 0 | 2 |
| HR output | 0 | 19 | 13 | 22 | 87 | 28 | 2 | 3 |
| HR filtered | 0 | 19 | 9 | 22 | 59 | 26 | 0 | 2 |

^a^ AR=Arkansas, USA; IA=Iowa, USA; MB=Manitoba, Canada; MI=Michigan, USA; MN=Minnesota, USA; OH=Ohio, USA; ON=Ontario, Canada; and WI=Wisconsin, USA.

Table S3. We developed home range (HR) estimates from trumpeter swan GPS collar data by individual for each breeding season that the collar transmitted data. Home range estimates were analyzed to better understand space use and were filtered to develop landscape suitability models with swan data from known breeders. We provide sample sizes per year and the number of swans with *n* years of data for all collared individuals, for individuals whose data were used to develop home range estimates, and for breeding individuals whose data we used for developing landscape suitability models.

|  | Individuals in year *t* | | | | |  | Number of Individuals with *n* year(s) of data | | | |
| --- | --- | --- | --- | --- | --- | --- | --- | --- | --- | --- |
| Group | 2019 | 2020 | 2021 | 2022 | 2023 |  | 1 year | 2 years | 3 years | 4 years |
| Collar data | 16 | 70 | 66 | 51 | 17 |  | 22 | 30 | 26 | 15 |
| HR output | 8 | 57 | 61 | 38 | 10 |  | 27 | 37 | 15 | 7 |
| HR filtered | 7 | 42 | 48 | 32 | 8 |  | 16 | 27 | 13 | 7 |

Table S4. We developed home range (HR) estimates from trumpeter swan GPS collar data by individual for each breeding season that the collar transmitted data, and for up to three periods within a season (early, middle, or late) if the data permitted. Home range estimates were analyzed to better understand space use over time and were filtered to develop landscape suitability models with swan data from known breeders with preference for the middle period, then early period, and then late period depending on the number of home range outputs available. We provide sample sizes for the number of home range estimates for each period across all years, and the number of swan-years with 1, 2, or 3 home range estimates in a year.

|  | Number of individual:year home range periods | | |  | Individual:year with *n* home range(s) periods | | |
| --- | --- | --- | --- | --- | --- | --- | --- |
| Group | Early | Middle | Late |  | 1 HR | 2HR | 3HR |
| HR output | 103 | 95 | 158 |  | 80 | 6 | 88 |
| HR breeder | 88 | 80 | 123 |  | 58 | 4 | 75 |
| HR in SDM | 9 | 80 | 48 |  | 137 |  |  |

Table S5. Pairwise comparisons (ratios) of estimated marginal means from 2 models that related trumpeter swan home-range (HR) or core-area (CA) sizes to breeding season period (early, middle, or late), sex (females or male), and breeding status (paired but breeding status unknown, or known breeder).

| Model | Term comparisons | Ratio (95% CI) | SE | *Z* | *P* |
| --- | --- | --- | --- | --- | --- |
| CA | Early/Late | 0.56 (0.39-0.82) | 0.09 | -3.55 | 0.001 |
| CA | Early/Middle | 0.74 (0.52-1.06) | 0.11 | -1.97 | 0.12 |
| CA | Late/Middle | 1.32 (0.95-1.85) | 0.19 | 1.97 | 0.12 |
| CA | Female/Male | 0.76 (0.37–1.57) | 0.28 | -0.736 | 0.46 |
| CA | Breeding/Paired | 0.39 (0.17-0.91) | 0.17 | -2.19 | 0.03 |
| HR | Early/Late | 0.82 (0.58–1.16) | 0.12 | -1.37 | 0.36 |
| HR | Early/Middle | 0.90 (0.65–1.24) | 0.12 | -0.80 | 0.70 |
| HR | Late/Middle | 1.10 (0.80–1.50) | 0.15 | 0.69 | 0.77 |
| HR | Female/Male | 0.77 (0.38–1.55) | 0.27 | -0.74 | 0.46 |
| HR | Breeding/Paired | 0.37 (0.17–0.84) | 0.15 | -2.40 | 0.02 |

Table S6. Estimated marginal means from 2 models that related trumpeter swan home-range (HR) or core-area (CA) sizes to breeding season period (early, middle, or late), sex (females or male), and breeding status (paired but breeding status unknown, or known breeder).

| Model | Term | Mean (95% CI) | SE |
| --- | --- | --- | --- |
| CA | Early | 47.3 (29.1-76.9) | 11.7 |
| CA | Middle | 63.6 (39.6-102.1) | 15.4 |
| CA | Late | 84.1 (53.9-131.3) | 19.1 |
| CA | Female | 55.2 (33.5–91.1) | 14.1 |
| CA | Male | 72.5 (38.7–135.6) | 23.2 |
| CA | Breeding | 39.6 (26.2–59.8) | 8.3 |
| CA | Paired | 101.0 (47.7–213.6) | 38.6 |
| HR | Early | 377.2 (237.9–598.2) | 88.7 |
| HR | Middle | 421.5 (267.9–663.3) | 97.5 |
| HR | Late | 462.1 (301.5-708.4) | 100.7 |
| HR | Female | 367.6 (227.5–593.8) | 90.0 |
| HR | Male | 477.3 (261.3–871.5) | 146.6 |
| HR | Breeding | 256.0 (172.4–380.1) | 51.6 |
| HR | Paired | 658.3 (334.0–1406.0) | 251.3 |

Table S7. Two model summaries where trumpeter swan home-range (HR) or core-area (CA) sizes were related to breeding season period (early, middle, or late), sex (females or male), and breeding status (paired but breeding status unknown, or known breeder). Reference levels include breeders, females, and the early breeding season period.

| Model | Term | β (95% CI) | SE | Z | *P* |
| --- | --- | --- | --- | --- | --- |
| CA | Intercept | 3.25 (2.71 – 3.79) | 0.28 | 11.77 | <0.001 |
| CA | Middle breeding season | 0.30 (0.002 - 0.59) | 0.15 | 1.97 | 0.05 |
| CA | Late breeding season | 0.58 (0.26 – 0.89) | 0.16 | 3.55 | <0.001 |
| CA | Male | 0.27 (-0.45 – 1.00) | 0.37 | 0.74 | 0.46 |
| CA | Paired | 0.94 (0.10 - 1.77) | 0.43 | 2.19 | 0.03 |
| HR | Intercept | 5.31 (4.80 – 5.84) | 0.26 | 20.23 | <0.001 |
| HR | Middle breeding season | 0.11 (-0.16 – 0.38) | 0.14 | 0.80 | 0.42 |
| HR | Late breeding season | 0.20 (-0.09 – 0.49) | 0.15 | 1.37 | 0.17 |
| HR | Male | 0.26 (-0.44 – 0.96) | 0.36 | 0.74 | 0.46 |
| HR | Paired | 0.98 (0.18 – 1.79) | 0.41 | 2.40 | 0.02 |

Table S8. Pairwise comparisons (ratios) of estimated marginal means from 2 models that related the distances between trumpeter swan home-range (HR) or core-area (CA) centroids across annual breeding season periods to breeding season periods (early to middle, middle to late, or early to late), sex (females or male), and breeding status (paired but breeding status unknown, or known breeder).

| Model | Pairs | Ratio | SE | *Z* | *P* |
| --- | --- | --- | --- | --- | --- |
| CA | Early to late/early to middle | 1.88 (1.60 – 2.21) | 0.13 | 9.06 | <0.001 |
| CA | Early to late/middle to late | 1.98 (1.68 – 2.34) | 0.14 | 9.72 | <0.001 |
| CA | Early to middle/middle to late | 1.05 (0.89 – 1.25) | 0.08 | 0.73 | 0.74 |
| CA | Female/Male | 0.97 (0.52 – 1.78) | 0.30 | -0.11 | 0.91 |
| CA | Breeding/Paired | 0.88 (0.39 – 1.99) | 0.37 | -0.31 | 0.75 |
| HR | Early to late/early to middle | 1.65 (1.38 – 1.97) | 0.13 | 6.47 | <0.001 |
| HR | Early to late/middle to late | 2.22 (1.84 – 2.66) | 0.17 | 10.15 | <0.001 |
| HR | Early to middle/middle to late | 1.34 (1.11 ­– 1.63) | 0.11 | 3.66 | <0.001 |
| HR | Female/Male | 0.57 (0.31 – 1.05) | 0.18 | -1.82 | 0.07 |
| HR | Breeding/Paired | 1.03 (0.46 – 2.33) | 0.43 | 0.08 | 0.94 |

Table S9. Estimated marginal means from 2 models that related the distances between trumpeter swan home-range (HR) or core-area (CA) centroids across annual breeding season periods to breeding season periods (early to middle, middle to late, or early to late), sex (females or male), and breeding status (paired but breeding status unknown, or known breeder).

| Model | Periods | Mean (95% CI) | SE |
| --- | --- | --- | --- |
| CA | Early to late | 350 (230 – 531) | 74.7 |
| CA | Early to middle | 186 (123 – 283) | 39.7 |
| CA | Middle to late | 177 (116 – 268) | 37.7 |
| CA | Female | 222 (137 – 361) | 55.0 |
| CA | Male | 229 (134 – 393) | 62.9 |
| CA | Breeding | 211 (151 – 295) | 36.0 |
| CA | Paired | 241 (114 – 510) | 92.1 |
| HR | Early to late | 401 (264 – 609) | 85.4 |
| HR | Early to middle | 243 (160 – 369) | 51.7 |
| HR | Middle to late | 181 (119 – 275) | 38.5 |
| HR | Female | 196 (121 – 318) | 48.4 |
| HR | Male | 345 (203 – 589) | 94.0 |
| HR | Breeding | 265 (190 – 369) | 44.8 |
| HR | Paired | 256 (122 – 538) | 97.0 |

Table S10. Two model summaries where the distances between trumpeter swan home-range (HR) or core-area (CA) centroids across annual breeding season periods were related to breeding season periods (early to middle, middle to late, or early to late), sex (females or male), and breeding status (paired but breeding status unknown, or known breeder). Reference levels include breeders, females, and the early to late breeding season periods.

| Model | Term | β (95% CI) | SE | Z | *P* |
| --- | --- | --- | --- | --- | --- |
| CA | Intercept | 5.77 (5.32 – 6.22) | 0.23 | 25.38 | <0.001 |
| CA | Early to middle period | -0.63 (-0.77 – -0.49) | 0.07 | -9.06 | <0.001 |
| CA | Middle to late period | -0.68 (-0.82 – -0.55) | 0.07 | -9.72 | <0.001 |
| CA | Male | 0.03 (-0.58 – 0.65) | 0.31 | 0.12 | 0.92 |
| CA | Paired | 0.13 (-0.69 – 0.95) | 0.42 | 0.31 | 0.75 |
| HR | Intercept | 5.73 (5.28 – 6.18) | 0.23 | 25.11 | <0.001 |
| HR | Early to middle period | -0.50 (-0.65 – -0.35) | 0.08 | -6.47 | <0.001 |
| HR | Middle to late period | -0.80 (-0.95 – -0.64) | 0.08 | -10.15 | <0.001 |
| HR | Male | 0.56 (-0.04 – 1.17) | 0.31 | 1.82 | 0.07 |
| HR | Paired | -0.03 (-0.85 – 0.78) | 0.41 | -0.08 | 0.94 |

Table S11. We developed home range (HR; 95% utilization distribution isopleth) and core area (CA; 50^th^ utilization distribution isopleth) estimates from trumpeter swan GPS collar data by individual for each breeding season that the collar transmitted data and for up to 3 periods within a season (early, middle, or late) if the data permitted. We provide summary statistics of core area (CA) and home range (HR) size and summary statistics grouped by breeding season period, sex, and breeding status.

| Area | Grouping factor | Factor level | Min. (ha) | Max. (ha) | Mean (ha) | Median (ha) | SD (ha) |
| --- | --- | --- | --- | --- | --- | --- | --- |
| CA | None | None | 0.16 | 12,256.22 | 367.30 | 16.01 | 1,245.61 |
| HR | None | None | 1.01 | 46,341.61 | 1,916.25 | 118.50 | 5,687.75 |
| CA | Period | Early | 0.51 | 7,418.53 | 321.45 | 8.59 | 1,067.81 |
| CA | Period | Late | 0.16 | 12,256.22 | 386.37 | 26.60 | 1,377.90 |
| CA | Period | Middle | 1.23 | 9,012.02 | 385.31 | 19.29 | 1,203.33 |
| HR | Period | Early | 7.25 | 30,932.88 | 1,726.95 | 90.11 | 5,327.53 |
| HR | Period | Late | 1.01 | 46,341.61 | 1,897.52 | 135.33 | 5,712.79 |
| HR | Period | Middle | 9.09 | 42,602.70 | 2,152.64 | 174.36 | 6,063.94 |
| CA | Breeding status | Breeder | 0.16 | 12,256.22 | 397.40 | 13.69 | 1,365.40 |
| CA | Breeding status | Paired | 1.25 | 1,833.55 | 232.57 | 65.59 | 368.78 |
| HR | Breeding status | Breeder | 1.01 | 46,341.61 | 2,035.87 | 98.73 | 6,214.18 |
| HR | Breeding status | Paired | 17.58 | 9,323.36 | 1,380.70 | 318.68 | 2,027.09 |
| CA | Sex | Female | 0.75 | 9,012.02 | 272.35 | 17.34 | 1,045.67 |
| CA | Sex | Male | 0.16 | 12,256.22 | 500.75 | 13.77 | 1,475.31 |
| HR | Sex | Female | 10.43 | 42,602.70 | 1,406.65 | 131.00 | 4,707.98 |
| HR | Sex | Male | 1.01 | 46,341.61 | 2,632.43 | 107.59 | 6,784.50 |

Table S12. We developed landscape suitability models by relating trumpeter swan occurrence and pseudo-absence data to local and landscape-scale summaries of the proportion of wetland and upland characteristics. These covariates were scaled by subtracting the mean and dividing by the standard deviation. We provide covariate summary statistics used for scaling for repeatability.

| Variable | Mean | SD |
| --- | --- | --- |
| High permanence flooded vegetation (590-m) | 0.047 | 0.116 |
| Grass (220-m) | 0.161 | 0.228 |
| Wetland perimeter (590-m) | 0.056 | 0.063 |
| High permanence open water (50-m) | 0.210 | 0.372 |
| Low permanence open water (50-m) | 0.052 | 0.149 |

Table S13. We developed landscape suitability models by relating trumpeter swan occurrence and pseudo-absence data to local and landscape-scale summaries of the proportion of wetland and upland characteristics. We used an exploratory multi-model comparison for model selection. For each term from the top-performing model we provide coefficient estimates, standard error, z-value, and p-value.

| Term | β (95% CI) | SE | Z | *P* |
| --- | --- | --- | --- | --- |
| Intercept | 0.23 (-0.12 – 0.57) | 0.18 | 1.30 | 0.195 |
| High permanence flooded vegetation (590-m) | 1.69 (1.59 – 1.80) | 0.05 | 30.98 | <0.001 |
| Grass (220-m) | 0.38 (0.33 – 0.42) | 0.02 | 15.32 | <0.001 |
| Wetland perimeter (590-m) | 1.14 (1.08 – 1.20) | 0.03 | 37.60 | <0.001 |
| High permanence open water (50-m) | 0.86 (0.81 – 0.91) | 0.03 | 32.90 | <0.001 |
| Low permanence open water (50-m) | 0.60 (0.52 – 0.69) | 0.04 | 14.40 | <0.001 |

Table S14. We developed landscape suitability models by relating trumpeter swan occurrence and pseudo-absence data to local and landscape-scale summaries of the proportion of wetland and upland characteristics. We used an exploratory multi-model comparison for model selection and after examining residuals versus fitted plots we explored covariate interactions and quadratics that reduce patterns. We compare the AICc and conditional and marginal R^2^ from the original model and for each quadratic and interaction that was tested.

| Model Update | AICc | ΔAICc from original model | Conditional R^2^ | Marginal R^2^ |
| --- | --- | --- | --- | --- |
| None (original) | 14,142.27 | 0.00 | 0.74 | 0.58 |
| Low permanence open water (50-m):grass (590-m) | 14,142.97 | 0.70 | 0.74 | 0.58 |
| Wetland perimeter (590-m):grass (220-m) | 14,142.20 | -0.08 | 0.74 | 0.58 |
| Grass (220-m)^2^ | 14,136.72 | -5.55 | 0.74 | 0.58 |
| High permanence open water (50-m):wetland Perimeter (590-m) | 14,136.30 | -5.98 | 0.74 | 0.58 |
| Low permanence open water (50-m):high permanence flooded vegetation (590-m) | 14,122.77 | -19.50 | 0.73 | 0.57 |
| High permanence open water (50-m):high permanence flooded vegetation (590-m) | 14,116.61 | -25.66 | 0.74 | 0.59 |
| High permanence open water (50-m):grass (220-m) | 14,116.40 | -25.87 | 0.74 | 0.58 |
| High permanence flooded vegetation (590-m)^2^ | 14,110.28 | -31.99 | 0.74 | 0.57 |
| High permanence open water (50-m):low permanence open water (50-m) | 14,100.84 | -41.43 | 0.74 | 0.58 |
| High permanence open water (50-m)^2^ | 14,056.24 | -86.03 | 0.74 | 0.58 |
| High permanence flooded vegetation (590-m):grass (220-m) | 14,035.96 | -106.31 | 0.74 | 0.58 |
| Low permanence open water (50-m):wetland perimeter (220-m) | 13,993.94 | -148.33 | 0.74 | 0.59 |
| Low permanence open water (50-m)^2^ | 13,991.54 | -150.74 | 0.74 | 0.58 |
| High permanence flooded vegetation (590-m):wetland perimeter (590-m) | 13,934.87 | -207.40 | 0.75 | 0.61 |
| Wetland perimeter (590-m)^2^ | 13,645.76 | -496.51 | 0.73 | 0.58 |
| Wetland perimeter (590-m)^2^+ wetland perimeter (590-m):high permanence flooded vegetation (590-m) + wetland perimeter (590-m)^2^:high permanence flooded vegetation (590-m) | 13,362.98 | -779.30 | 0.77 | 0.63 |

Table S15. Trumpeter swan range expansion model summary that related total range area (sum of geographical units occupied) in the western Great Lakes region of North America to year and survey intensity. We considered geographical units (50-km hex grids) part of the trumpeter swan range once it had 2 annual occurrence records within a 6-year period. Occurrence data were obtained from long-term structured (i.e., The North American Breeding Bird Survey) and semi-structured citizen science datasets (i.e., eBird and iNaturalist).

| Term | β (95% CI) | SE | *t* | *P* |
| --- | --- | --- | --- | --- |
| Intercept | 11.95 (11.58 – 12.32) | 0.19 | 62.83 | <0.001 |
| Year | 2.51 (2.05 – 2.96) | 0.23 | 10.81 | <0.001 |
| Year^2^ | -0.25 (-0.31 – -0.19) | 0.03 | -8.13 | <0.001 |
| Log Mean Effort | 0.18 (0.08 – 0.28) | 0.05 | 3.64 | 0.002 |

Table S16. Trumpeter swan range-expansion model summary that related newly occupied geographical units and unoccupied units to suitable conditions within those units, distance to previously occupied units, year, and survey effort. We considered geographical units (50-km hex grids) part of the trumpeter swan range once it had ≥2 annual occurrence records within a 6-year period. Occurrence data were obtained from long-term structured (i.e., The North American Breeding Bird Survey) and semi-structured citizen science datasets (i.e., eBird and iNaturalist).

| Term | β (95% CI) | SE | *Z* | *P* |
| --- | --- | --- | --- | --- |
| Intercept | 233.86 (145.15 – 326.21) | 46.12 | 5.07 | <0.001 |
| Distance | -0.02 (-0.03 – -0.02) | 0.002 | -8.91 | <0.001 |
| Year | -0.12 (-0.16 – -0.07) | 0.02 | -5.06 | <0.001 |
| Landscape Suitability | 7.88 (-0.45 – 14.87) | 4.03 | 1.96 | 0.05 |
| Log Mean Effort | 0.53 (0.40 – 0.67) | 0.07 | 7.94 | <0.001 |

Figure S1. Capture locations (black points) of Interior Population Trumpeter Swans collared with GPS-GSM transmitters from July 2019–January 2022.

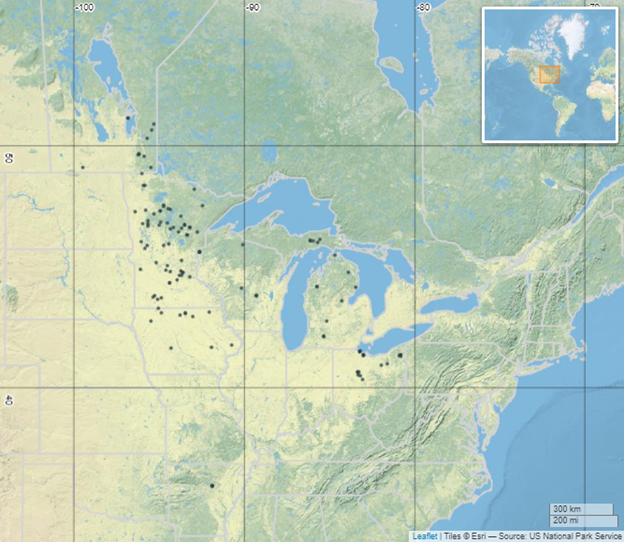

Figure S2. Marginal effects plots of the top performing trumpeter swan landscape suitability model from an exploratory multi-model comparison. Shaded areas are 95% confidence intervals. The x-axis value labels include the scaled value (standard deviations from the mean) and their respective landscape-scale proportions.

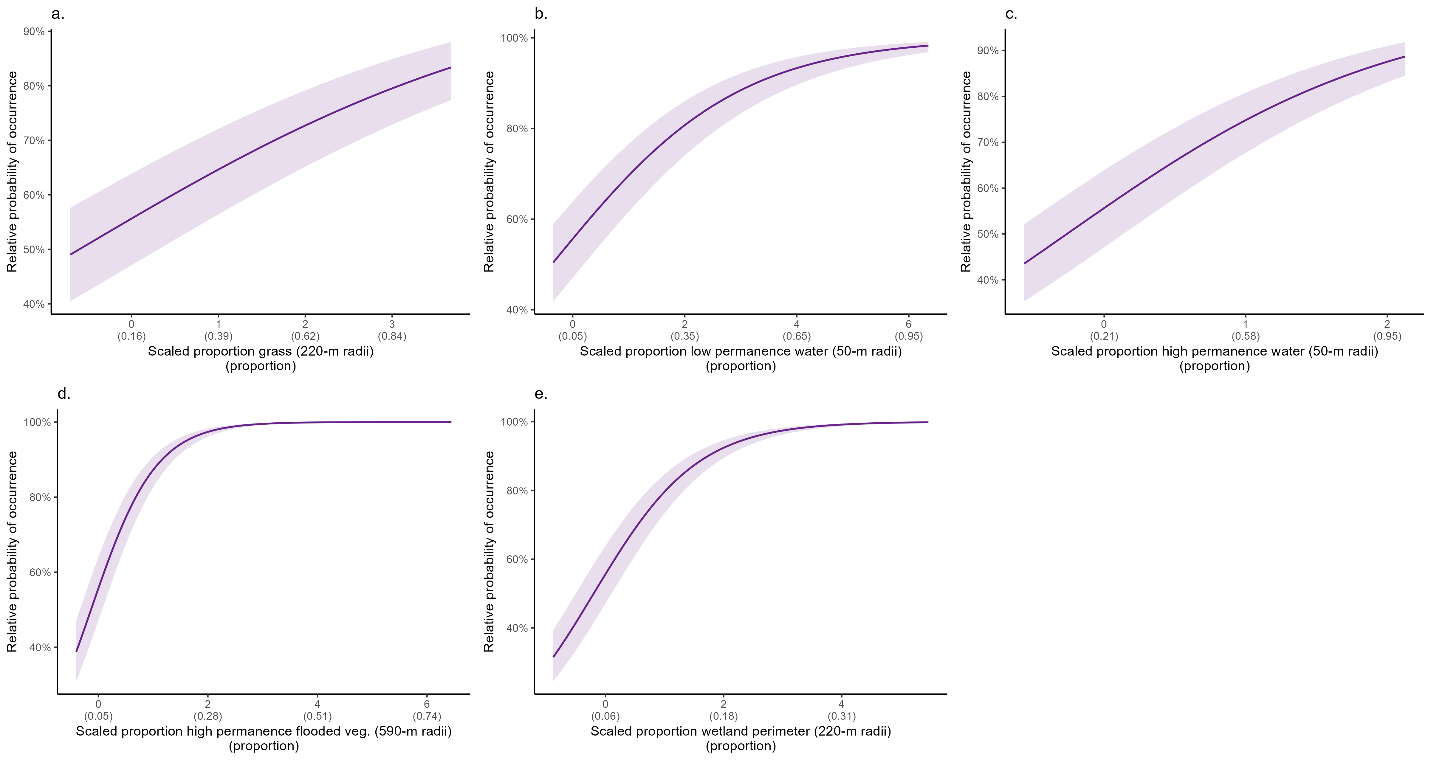

Figure S3. eBird survey intensity (2004-2023) represented as the number of checklists submitted within 50-km hex grids in the western Great Lakes region of North America. These data were used in trumpeter swan range expansion models to account for spatial and temporal bias of survey effort. The line graph represents mean annual survey intensity (effort) across all grid cells.

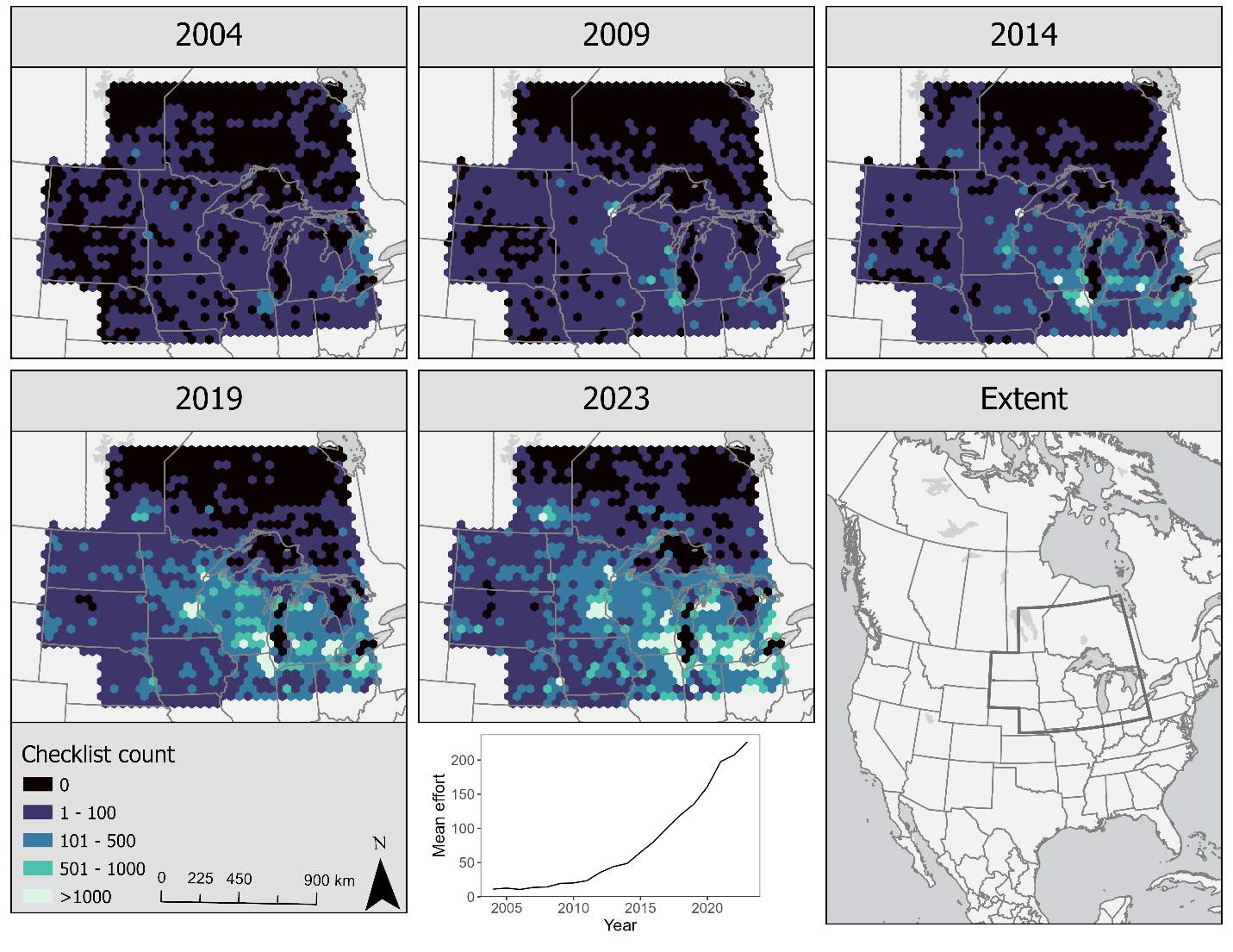
